# Loss of mitochondrial pyruvate transport initiates cardiac glycogen accumulation and heart failure

**DOI:** 10.1101/2024.06.06.597841

**Authors:** Rachel C. Weiss, Kelly D. Pyles, Kevin Cho, Michelle Brennan, Jonathan S. Fisher, Gary J. Patti, Kyle S. McCommis

**Affiliations:** Edward A. Doisy Department of Biochemistry & Molecular Biology, Saint Louis University School of Medicine; Departments of Chemistry, Medicine, and Center for Mass Spectrometry and Metabolic Tracing, Washington University in St. Louis; Department of Biology, Saint Louis University

**Author notes:** For correspondence: Kyle S. McCommis, PhD 1100 South Grand Blvd. St. Louis, MO 63104. K.S.M. maintained full access to all study data and takes responsibility for its integrity and that of the data analyses.

**Keywords:** heart failure, mitochondria, pyruvate, ketogenic diet

## Abstract

Heart failure involves metabolic alterations including increased glycolysis despite unchanged or decreased glucose oxidation. The mitochondrial pyruvate carrier (MPC) regulates pyruvate entry into the mitochondrial matrix, and cardiac deletion of the MPC in mice causes heart failure. How MPC deletion results in heart failure is unknown. We performed targeted metabolomics and isotope tracing in wildtype (fl/fl) and cardiac-specific Mpc2-/- (CS-Mpc2-/-) hearts after in vivo injection of U-^13^C-glucose. Failing CS-Mpc2-/- hearts contained normal levels of ATP and phosphocreatine, yet these hearts displayed increased enrichment from U-^13^C-glucose and increased glycolytic metabolite pool sizes. ^13^C enrichment and pool size was also increased for the glycogen intermediate UDP-glucose, as well as increased enrichment of the glycogen pool. Glycogen levels were increased ∼6-fold in the failing CS-Mpc2-/- hearts, and glycogen granules were easily detected by electron microscopy. In young, non-failing CS-Mpc2-/- hearts, increased glycolytic ^13^C enrichment occurred, but glycogen levels remained low and unchanged compared to fl/fl hearts. Inhibiting glycogen synthase with MZ-101 reduced cardiac glycogen levels and improved heart failure. Feeding a ketogenic diet to CS-Mpc2-/- mice reversed the heart failure and normalized the cardiac glycogen and glycolytic metabolite accumulation. Cardiac glycogen levels were also elevated in mice infused with angiotensin-II, and both the cardiac hypertrophy and glycogen levels were improved by ketogenic diet. Thus, loss of MPC in the heart causes glycogen accumulation and heart failure, while inhibition of glycogen synthesis or a ketogenic diet can reverse both the glycogen accumulation and heart failure.

## Introduction

Currently, 6.7 million adults in the United States suffer from heart failure, which has risen from 6.0 million in just two years^1^. Those diagnosed with heart failure have a 5-year mortality rate of over 50%, which has remained unchanged in recent years despite existing treatment options^1^. Heart failure can arise from a multitude of causes, including ischemia/infarction, cardiac hypertrophy, chronic hypertension, or valvular diseases^1,2^. A better understanding of the development and pathophysiology of heart failure is needed to generate improved treatment options.

Since the heart lacks the ability to store ample amounts of metabolites or high energy phosphates, a continuous supply of nutrients is required to fuel its constant contractile work. Mitochondrial oxidative metabolism plays a large role in the production of high energy phosphates in the adult heart. The preferred fuel source of the heart are fatty acids, with fatty acid oxidation (FAO) accounting for ∼70% of the adenosine triphosphate (ATP) produced^3,4^. Yet the healthy heart is metabolically flexible and can adapt to use alternative fuel sources to meet its energy demands, including the use of glucose, lactate, ketone bodies, and amino acids^3,4^. Glycolysis accounts for ∼5% of ATP produced in the healthy heart^3,4^. However, during hypertrophy and heart failure metabolic changes occur, including significantly reduced FAO, increased glycolysis, but relatively decreased glucose oxidation^3–6^. This mismatch between increased glycolysis, yet unchanged or decreased glucose oxidation has been suggested to be important for stimulating cardiac hypertrophy and failure^7–9^.

The mitochondrial pyruvate carrier (MPC) is responsible for transporting pyruvate, the end product of glycolysis or oxidized lactate, into the mitochondrial matrix^10,11^. The mammalian MPC is a heterodimeric complex composed of MPC1 and MPC2 proteins, both of which are necessary for MPC complex formation and pyruvate transport^10,11^. Once pyruvate has been transported into the mitochondria, it can be converted into acetyl-CoA and used in the citric acid cycle^10,11^. Recent studies by us and others revealed that cardiac MPC gene and protein expression is significantly reduced in humans with heart failure^12–15^ and that deletion of MPC1^4,12,14^ or MPC2^12,13^ in mice leads to spontaneous heart failure with age. However, it is still not understood how reduction or deletion of MPC expression contributes to heart failure.

In the present study, we suggest that this heart failure is associated with accumulation of glycogen in the MPC-deficient heart. Even with dilated cardiomyopathy, cardiac-specific MPC2-/- (CS-Mpc2-/-) hearts contain normal levels of ATP and phosphocreatine compared to wildtype littermates, suggesting that lack of pyruvate oxidation does not simply result in “energetic stress”. Glucose uptake is increased, and glycolytic metabolites accumulate in both young, non-failing and failing CS-Mpc2-/- hearts. However, glycogen accumulation is only observed in failing CS-Mpc2-/- hearts and the heart failure and glycogen accumulation can be improved by inhibiting glycogen synthase or providing a high-fat, low-carbohydrate ketogenic diet (KD). Lastly, we also observe increased glycogen in hearts from angiotensin-II (AngII) infused mice, a hypertrophic stimulus that reduces *Mpc* expression^12^, and both the cardiac glycogen and hypertrophy in this model are also normalized by KD feeding.

## Results

### Heart failure in CS-Mpc2-/- mice is not due to energetic stress

We and others have previously described spontaneous heart failure development in MPC-deficient murine hearts^4,12–14^, but what causes these hearts to fail is not understood. In attempt to uncover insights we performed bulk RNA sequencing on CS-Mpc2-/- hearts with dilated cardiomyopathy and age-matched fl/fl littermates (Figs. 1a-c). This analysis uncovered 956 significantly differentially expressed genes in the failing hearts compared to control, with ribosome and protein synthesis pathways upregulated and multiple pathways related to fatty acid metabolism, pyruvate metabolism, and amino acid catabolism downregulated (Fig. 1c). The downregulation of fatty acid metabolism is a well-known metabolic alteration in failing hearts^2^, including in these CS-Mpc2-/- hearts^13^. Despite transcriptional downregulation of fatty acid, pyruvate, and amino acid metabolic pathways, surprisingly CS-Mpc2-/- hearts maintained normal levels of the high energy phosphates ATP and phosphocreatine as well as normal phosphocreatine/creatine and phosphocreatine/ATP ratios (Figs. 1d-j). Although AMP levels were slightly elevated and ADP levels trended to be elevated, the lack of a decrease in ATP or phosphocreatine suggests normal energetic status of these failing CS-Mpc2-/- hearts. Additionally, there were no significant differences in nicotinamide nucleotides NAD+, NADH, NADP+, or NADP in failing CS-Mpc2-/- hearts (Figs. 1k-n), suggesting that redox control of metabolic pathways is also normal. Altogether, these results suggest that despite multiple decreased oxidative pathways, the heart failure in CS-Mpc2-/- mice is not caused by overt bioenergetic stress.

**Fig. 1:**
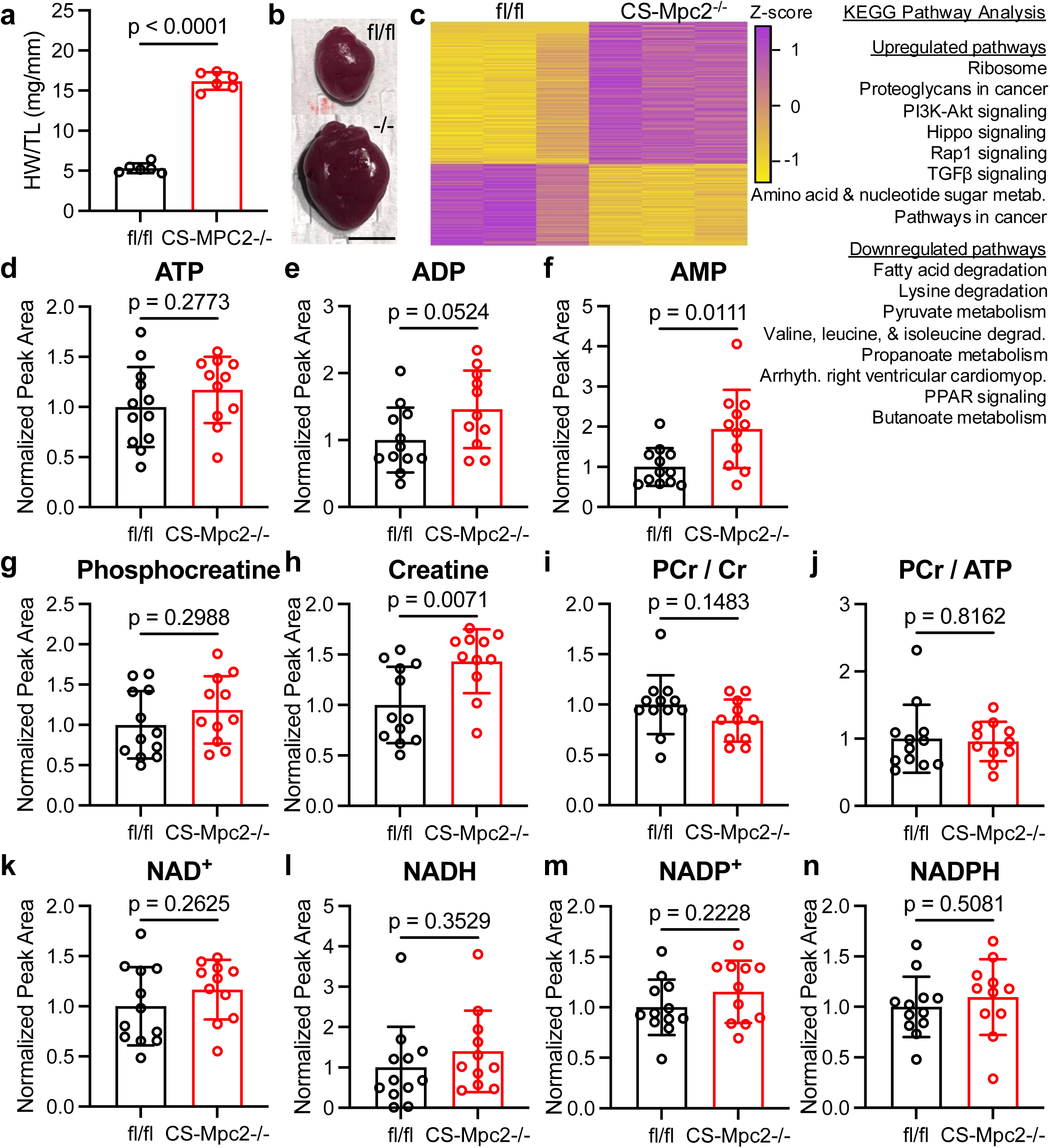
Heart Failure in CS-Mpc2-/- mice is not due to energetic stress. **a** Heart weight/tibia length (HW/TL) in CS-Mpc2-/- and fl/fl littermates, n=6. **b** Representative gross images of excised fl/fl and CS-Mpc2-/- hearts. Scale bar = 5 mm. **c** Heatmap of differentially expressed genes identified by RNA sequencing of hearts from fl/fl and CS-Mpc2-/- littermates, n=3. Most prominently dysregulated KEGG pathways upregulated and downregulated are listed. **d-n** mass spectrometry normalized peak area pool sizes of adenosine triphosphate (ATP) **(d)**, adenosine diphosphate (ADP) **(e)**, adenosine monophosphate (AMP) **(f)**, phosphocreatine **(g)**, creatine **(h)**, phosphocreatine/creatine (PCr/Cr) ratio **(i)**, PCr/ATP ratio **(j)** oxidized nicotinamide adenine dinucleotide (NAD^+^) **(k)**, reduced nicotinamide adenine dinucleotide (NADH) **(l)**, oxidized nicotinamide adenine dinucleotide phosphate (NADP^+^) **(m)**, and reduced nicotinamide adenine dinucleotide phosphate (NADPH) **(n)** in hearts of fl/fl and CS-Mpc2-/- littermates that had been injected i.p. with U-^13^C-glucose, n=11-12. Data are presented as mean±SD. Data were evaluated by unpaired, two-tailed Student’s t-test with Welch correction.

### Failing CS-Mpc2-/- hearts display increased glycolysis

As it has been well described that heart failure is associated with increased glucose utilization, often with unchanged or decreased glucose oxidation^16^, we hypothesized that MPC deletion would mimic these metabolic changes. To evaluate glucose metabolism, we injected mice intraperitoneally (i.p.) with U-^13^C-glucose, excised the hearts 30 min post injection, and assessed metabolite pool sizes and ^13^C isotopologues by UHPLC-MS/MS. We first focused on fully ^13^C labeled glycolytic metabolites (Fig. 2a) and their pool sizes (Supplementary Fig. 1). While glucose enrichment and pool size were unaltered, glucose-6-phosphate (G6P) enrichment showed a trend to be increased and all other downstream glycolytic metabolites displayed significantly greater ^13^C enrichment and pool size in CS-Mpc2-/- hearts (Figs. 2b-g and Supplementary Figs. 1a-h). Additionally, CS-Mpc2-/- hearts displayed increased enrichment and pool sizes of both lactate and alanine (Figs. 2h-i and Supplementary Figs. 1i-j), indicative of decreased pyruvate carbon entry into the TCA cycle. Altogether, these results suggest increased glycolytic metabolism in failing CS-Mpc2-/- hearts.

**Fig. 2:**
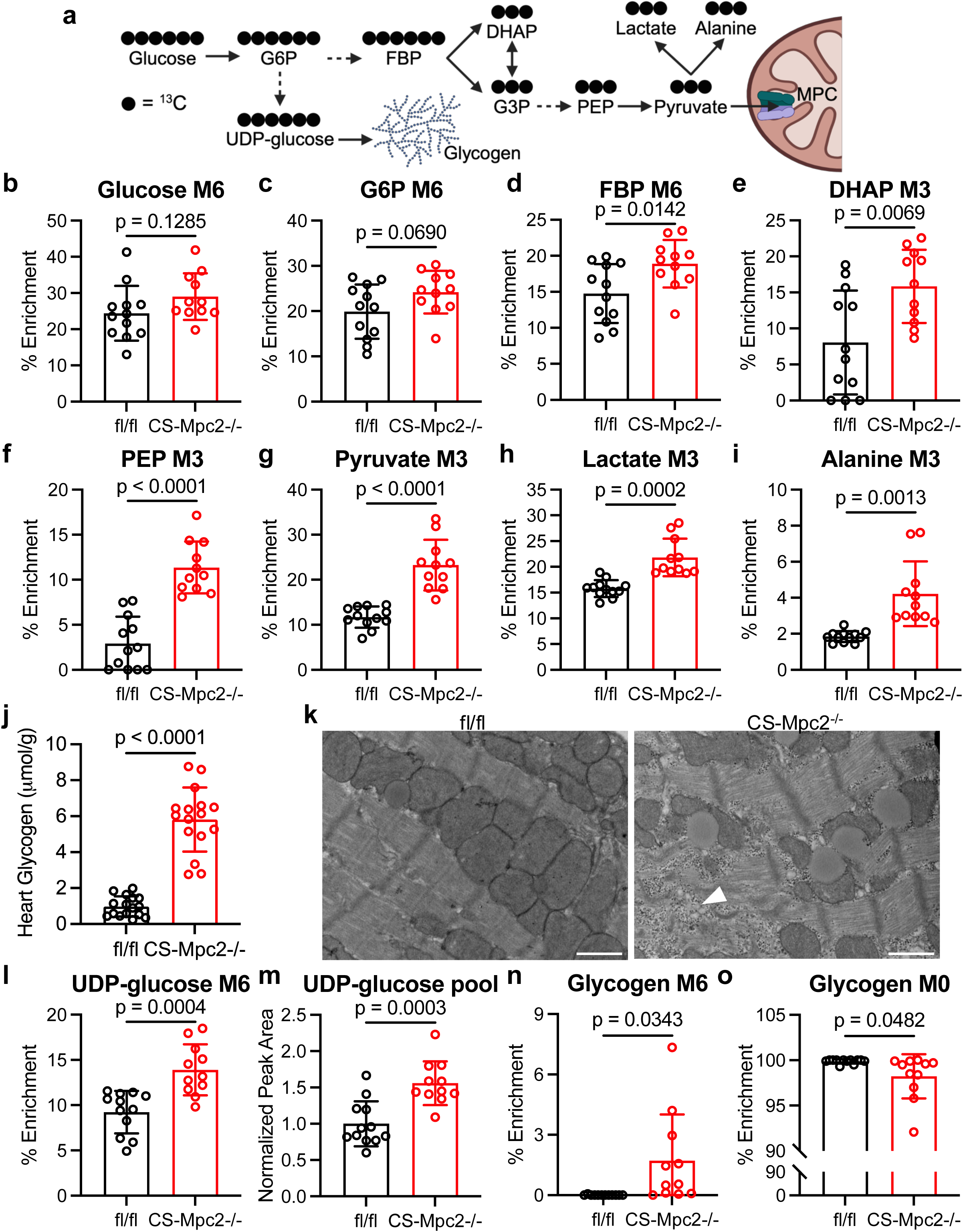
Failing CS-Mpc2-/- hearts display increased glycolysis and accumulate glycogen. **a** Schematic of U-^13^C-glucose labeling of glycolytic metabolites and glycogen, created with BioRender. **b-i** ^13^C % enrichment of glucose **(b)**, glucose-6-phosphate (G6P) **(c)**, fructose bisphosphate (FBP) **(d)**, dihydroxyacetone phosphate (DHAP) **(e)**, phosphoenolpyruvate (PEP) **(f)**, pyruvate **(g)**, lactate **(h),** and alanine **(i)** in hearts from fl/fl and CS-Mpc2-/- mice after injection with U-^13^C-glucose, n=11-12. **j** Cardiac glycogen levels measured biochemically in fl/fl and CS-Mpc2-/- littermates, n=16. **k** Representative transmission electron microscopy images at 5,000X magnification of the cardiac papillary muscle of fl/fl and CS-Mpc2-/-, white arrowhead denotes glycogen granules. Scale bars = 1 μm. **l-m** ^13^C % enrichment (**l**) and pool size (**m**) of uridine diphosphate (UDP)-glucose, n=11-12. **n-o** ^13^C % enrichment of the glucose within glycogen for fully labeled M6 **(n),** and fully unlabeled M0 **(o)**, n=11-12. Data are presented as mean±SD. Data were evaluated by unpaired, two-tailed Student’s t-test with Welch correction. See also Supplementary Figs. 1-4.

Surprisingly, CS-Mpc2-/- hearts do not display significantly decreased enrichment of TCA cycle metabolites (Supplementary Figs. 2a-l). Likewise, pool sizes of TCA cycle metabolites were also unaltered in CS-Mpc2-/- hearts apart from α-ketoglutarate being significantly decreased (Supplementary Figs. 2a-l). This normal TCA cycle enrichment from U-^13^C-glucose contrasts studies of MPC deletion or inhibition in other cell types, such as hepatocytes^17^, when provided U-^13^C-glucose or U-^13^C-pyruvate. However, while one previous study identified decreased TCA cycle enrichment with MPC1 deletion in isolated cardiomyocytes^14^, another previous study observed normal or even significantly increased enrichment of the TCA cycle in MPC1-/- hearts perfused with ^13^C-glucose^4^. It has recently been suggested that mitochondrial lactate import and metabolism could explain pyruvate carbon entry into MPC-/- cardiac mitochondria^18^. While the puzzling ^13^C enrichment results do not support it, it is expected and likely that MPC deletion reduces cardiac pyruvate oxidation, as both isolated mitochondria and permeabilized cardiac muscle fibers display significantly reduced oxygen consumption when provided with pyruvate/malate as substrates (Supplementary Fig. 3)^13^. Additionally, while mitochondria from fl/fl hearts are sensitive to the MPC inhibitor UK-5099 which decreases pyruvate respiration, there is little-to-no effect of UK-5099 in CS-Mpc2-/- mitochondria or muscle fibers (Supplementary Fig. 3). Potential explanations for these surprising TCA cycle ^13^C enrichment results are discussed in more detail below.

### CS-Mpc2-/- hearts do not increase glucose enrichment of the pentose phosphate pathway, but do increase flux into glycerol-3-phosphate

Several recent studies have suggested the importance of glucose flux through the pentose phosphate pathway to fuel hypertrophied or failing hearts with amino acids, RNA, and nucleotides^7,9^. Interestingly, in failing CS-Mpc2-/- hearts we observed no significant increases in ^13^C enrichment of 6-phosphogluconate, ribose-5-phosphate, sedoheptulose-7-phosphate, nor serine or glycine which can be end products of the pentose phosphate pathway (Supplementary Figs. 4a-f). As described above, we also identified no changes in NADP+ or NADPH (Figs. 1m-n), indicative of unaltered flux through the pentose phosphate pathway. Several pentose phosphate pathway metabolites displayed increased pool sizes in CS-Mpc2-/- hearts (Supplementary Figs. 4b-f), however the unaltered enrichment from U-^13^C glucose suggests that CS-Mpc2-/- hearts do not significantly increase glucose incorporation into the pentose phosphate pathway. Lastly, we observed increased ^13^C enrichment and pool size of glycerol-3-phosphate (Supplementary Figs. 1a,k-l), suggesting the failing CS-Mpc2-/- hearts increasingly shuttle glucose carbons into glycerophospholipid synthesis.

### Failing CS-Mpc2-/- hearts accumulate glycogen

Another potential fate of glucose is storage as glycogen, and indeed we^13^ and others^4^ previously observed increased glycogen content in MPC-deficient hearts, without expanding upon this finding. Here we confirmed again increased glycogen concentrations in failing CS-Mpc2-/- hearts measured biochemically (Fig. 2j) as well as by detection of abundant glycogen granules on transmission electron microscopy images of failing CS-Mpc2-/- hearts (Fig. 2k). In hearts from U-^13^C-glucose injected mice, we observed significantly elevated ^13^C enrichment and pool size of UDP-glucose (Figs. 2l-m), indicating increased flow of carbon into this glycogen intermediate. When glucose was liberated from the cardiac glycogen, fl/fl hearts displayed essentially zero ^13^C enrichment, while all but one of the CS-Mpc2-/- hearts displayed detectable enrichment (Fig. 2n). Conversely, the percentage of completely unlabeled glucose liberated from glycogen was significantly decreased in CS-Mpc2-/- hearts (Fig. 2o). Taken together, these results suggest that loss of MPC in the heart results in increased glycogen synthesis and accumulation of glycogen.

### Glucose uptake is increased, and glycogen synthase should be inhibited in CS-Mpc2-/- hearts

While the ^13^C-glucose enrichment studies suggest increased glycolysis in CS-Mpc2-/- hearts, the bolus of glucose injected makes it difficult to determine if cardiac glucose uptake is altered, and we observed similar labeling of glucose in fl/fl and CS-Mpc2-/- hearts in those studies (Fig. 2b). To more appropriately assess glucose uptake, left ventricle myocardial muscle fibers were incubated with ^3^H-2-deoxyglucose in the absence or presence of insulin. Basal glucose uptake in CS-Mpc2-/- myocardial fibers was nearly double that of fl/fl myocardial fibers and remained significantly higher with mild insulin-stimulation (Fig. 3a). Since heart failure is often associated with cardiac insulin-resistance^19^, we assessed insulin signaling by blotting for phosphorylated and total AKT after i.p. insulin injection, which was not significantly altered in CS-Mpc2-/- mice (Fig. 3b). Lastly, we also investigated the regulatory enzymes for glycogen synthesis and degradation. Interestingly, in the failing CS-Mpc2-/- hearts (Figs. 3c-e), we observed significantly elevated phosphorylation of glycogen synthase at serine 640 (Figs. 3c,f), which is a strong inhibitory phosphorylation site^20^. Additionally, expression of glycogenin-1, which acts as a primer for glycogen synthase by initializing the glucosyl chain^21^, was decreased in failing CS-Mpc2-/- hearts (Figs. 3c,g). We did not observe any difference in glycogen phosphorylase expression, therefore cannot suggest whether glycogenolysis, or the breakdown of glycogen, is altered in CS-Mpc2-/- hearts. Altogether, these results suggest that cardiac MPC loss increases glucose uptake and glycogen synthesis, despite the heart’s attempt to inhibit glycogen synthesis by phosphorylation of glycogen synthase and decreased glycogenin-1 expression.

**Fig. 3:**
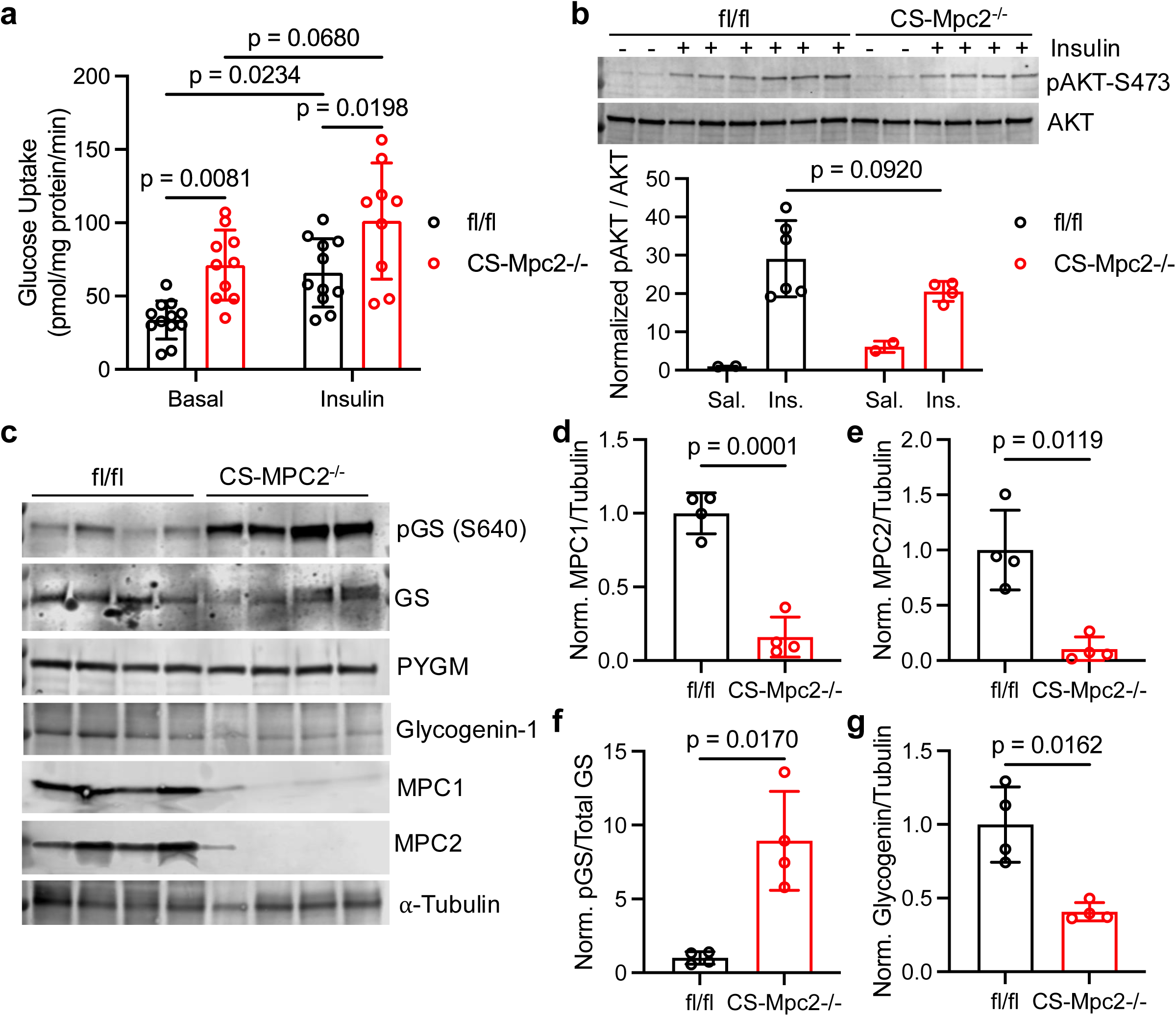
Glucose uptake is increased, and glycogen synthesis should be inhibited in CS-Mpc2-/- hearts. **a** Glucose uptake assessed by 2-deoxyglucose in the present or absence of insulin in cardiac muscle fibers from fl/fl and CS-Mpc2-/- littermates, n=9-12. **b** Western blot images of phosphorylated (serine-473) and total protein kinase B (AKT) and the ratio of quantified densitometry for pAKT/AKT from hearts of fl/fl and CS-Mpc2-/- mice injected with either saline vehicle or 5mU/g insulin. **c-g** Western blot images of phosphorylated glycogen synthase (pGS) serine-640, total glycogen synthase (GS), muscle glycogen phosphorylase (PYGM), glycogenin-1, mitochondrial pyruvate carrier (MPC) 1, MPC2, and α-tubulin (**c**), and the normalized densitometry quantification for MPC1/tubulin **(d)**, MPC2/tubulin **(e)**, pGS/total GS **(f)**, and glycogenin-1/tubulin **(g)** in hearts of fl/fl and CS-Mpc2-/- littermates, n=4. Data are presented as mean±SD. Data in **a** were evaluated by two-way analysis of variance (ANOVA) with Tukey post-hoc multiple-comparisons test. Data in **b** were evaluated by unpaired, two-tailed Student’s t-test with Welch correction due to only n=2 in saline groups. Data in **d-g** were evaluated by unpaired, two-tailed Student’s t-test with Welch correction.

### Non-failing CS-Mpc2-/- hearts from young mice display increased glycolysis, but no glycogen accumulation

As increased glucose uptake and glycolysis are known to occur in failing hearts, we investigated whether these metabolic changes were due to the presence of heart failure in CS-Mpc2-/- hearts, or a direct consequence of MPC deletion. To assess this, we repeated the U-^13^C-glucose enrichment experiment in 6-week-old fl/fl and CS-Mpc2-/- mice before the onset of cardiac remodeling and dysfunction^13^. The ^13^C enrichment of all glycolytic metabolites was increased in these young, non-failing CS-Mpc2-/- hearts (Figs. 4a-i). The pool size of glucose in these hearts was increased, but the pool size of all glycolytic metabolites was unaltered in the non-failing CS-Mpc2-/- hearts (Supplementary Figs. 5a-j). However, neither UDP-glucose enrichment nor pool size were significantly altered in the non-failing CS-Mpc2-/- hearts (Figs. 4j-k), suggesting no increase in glycogen synthesis. Indeed, glycogen concentrations were normal in these young, non-failing CS-Mpc2-/- hearts (Fig. 4l) and unlike in the failing hearts, we did not readily detect glycogen granules on electron micrographs (Fig. 4m). The increased glycolysis in these non-failing CS-Mpc2-/- hearts did not appear to increase pentose phosphate pathway flux as ribose-5-phosphate, and the enrichment of serine and glycine and their pool sizes remained normal (Supplementary Figs. 5k-o). However, enrichment into glycerol-3-phosphate was increased, while pool size remained unchanged (Supplementary Figs. 5p,q), suggesting increased flow of glucose carbons into glycerophospholipid synthesis. As within the failing CS-Mpc2-/- hearts, these nonfailing CS-Mpc2-/- hearts also displayed normal enrichment and pool size of TCA cycle intermediates after ^13^C-glucose injection (Supplementary Fig. 6). Cumulatively, these data suggest that cardiac MPC deletion, on its own, increases glycolytic metabolism, but only with the development of heart failure does glycogen accumulate.

**Fig. 4:**
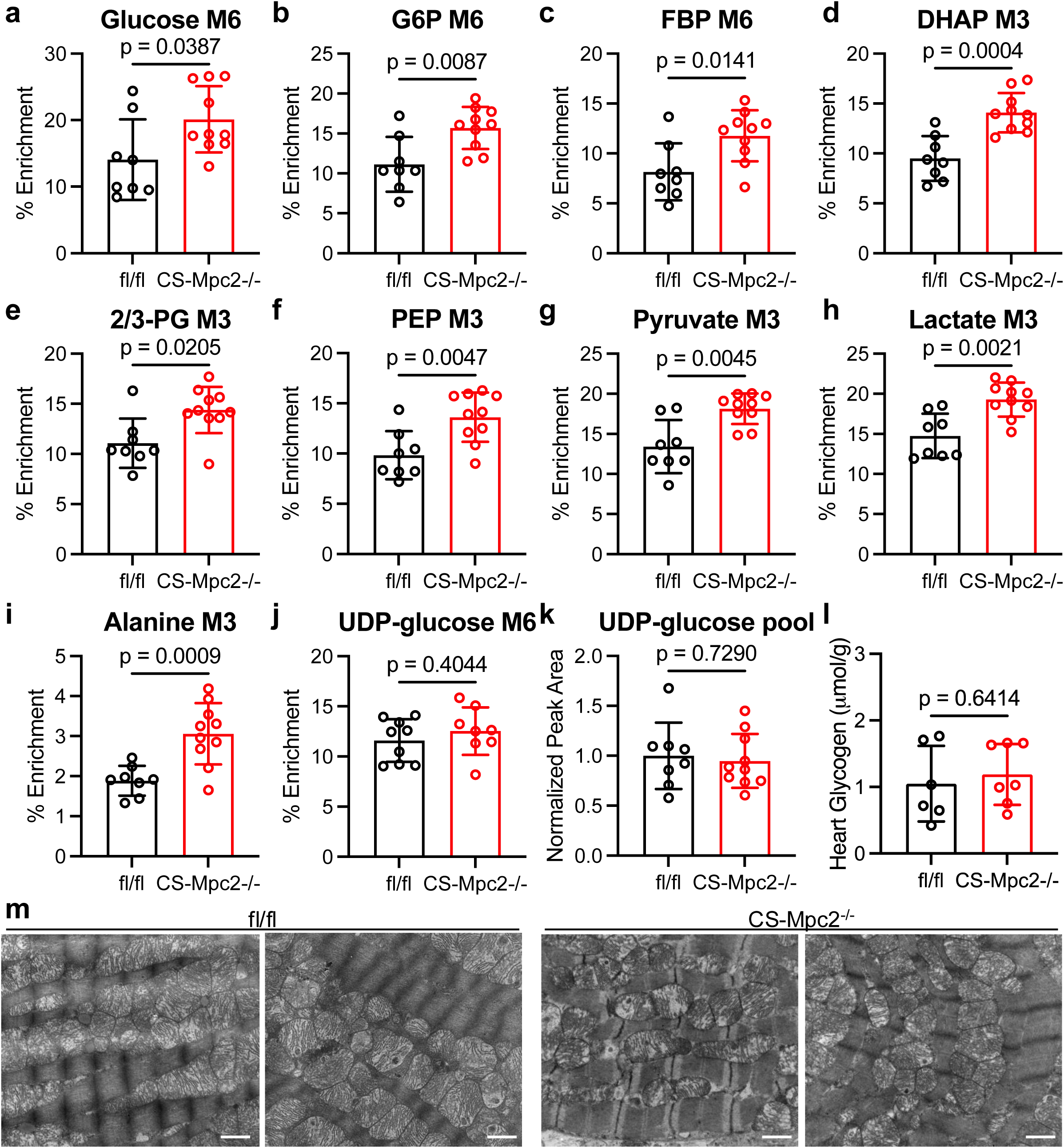
Non-failing CS-Mpc2-/- hearts from young mice display increased glycolysis, but no glycogen accumulation. a-k. ^13^C % enrichment of glucose **(a)**, gluocse-6-phosphate (G6P) **(b)**, fructose bisphosphate (FBP) **(c)**, dihydroxyacetone phosphate (DHAP) **(d)**, 2/3-phosphoglycerate (PG) **(e)**, phosphoenolpyruvate (PEP) **(f)**, pyruvate **(g)**, lactate **(h)**, alanine **(i)**, and uridine diphosphate (UDP)-glucose **(j)**, and the UDP-glucose normalized pool size (**k**) in hearts of young non-failing CS-Mpc2-/- and fl/fl littermates injected i.p. with U-^13^C-glucose, n=8-10. **l** Heart glycogen content measured biochemically in young non-failing CS-Mpc2-/- and fl/fl littermates, n=6-7. **m** Representative transmission electron microscopy images at 15,000X magnification of the cardiac papillary muscle in young fl/fl and CS-Mpc2-/- littermates. Scale bars = 1 μm. Data are presented as mean±SD, and were analyzed by unpaired, two-tailed Student’s t-test with Welch correction. See also Supplementary Figs. 5-6.

### Inhibition of glycogen synthase improves heart failure in CS-Mpc2-/- mice

To evaluate if glycogen accumulation had a causal relationship with heart failure development, we next attempted to decrease cardiac glycogen by treating mice with a recently developed glycogen synthase 1 (GYS1) inhibitor, MZ-101^22^. This potent small molecule inhibitor was recently shown to decreased glycogen accumulation in both skeletal muscle^22,23^ and the heart^23^ in mouse models of Pompe disease, a glycogen storage disease also associated with cardiomyopathy. MZ-101 treatment had no significant impact on blood glucose or lactate measurements (Figs. 5a,b). While vehicle-treated CS-Mpc2-/- mice displayed cardiac glycogen accumulation, glycogen levels were completely normalized in CS-Mpc2-/- mice treated with MZ-101 (Fig. 5c). Somewhat surprisingly, cardiac glycogen levels were unaffected in fl/fl mice treated with MZ-101, which was also observed previously in normal WT mice^23^, likely due to the low levels of glycogen in normal hearts. In slow twitch soleus skeletal muscle, glycogen content in both fl/fl and CS-Mpc2-/- mice was significantly decreased by MZ-101 (Fig. 5d), whereas there was no significant impact on liver glycogen content (Fig. 5e), since MZ-101 inhibits only GYS1 and not hepatic GYS2^22^. Dilated cardiomyopathy was significantly reduced as indicated by reduced heart weight and histology (Figs. 5f,g). Gene expression for natriuretic peptides, markers of cardiac failure, were elevated in vehicle-treated CS-Mpc2-/- hearts, and reduced by MZ-101 (Figs. 5h,i). Lastly, phosphorylation of glycogen synthase serine-640 was increased in vehicle-treated CS-Mpc2-/- failing hearts and was significantly decreased by MZ-101 (Fig. 5j). All together these data indicate that inhibition of glycogen synthase reduces glycogen accumulation leading to the improvement of heart failure in CS-Mpc2-/- mice.

**Fig. 5:**
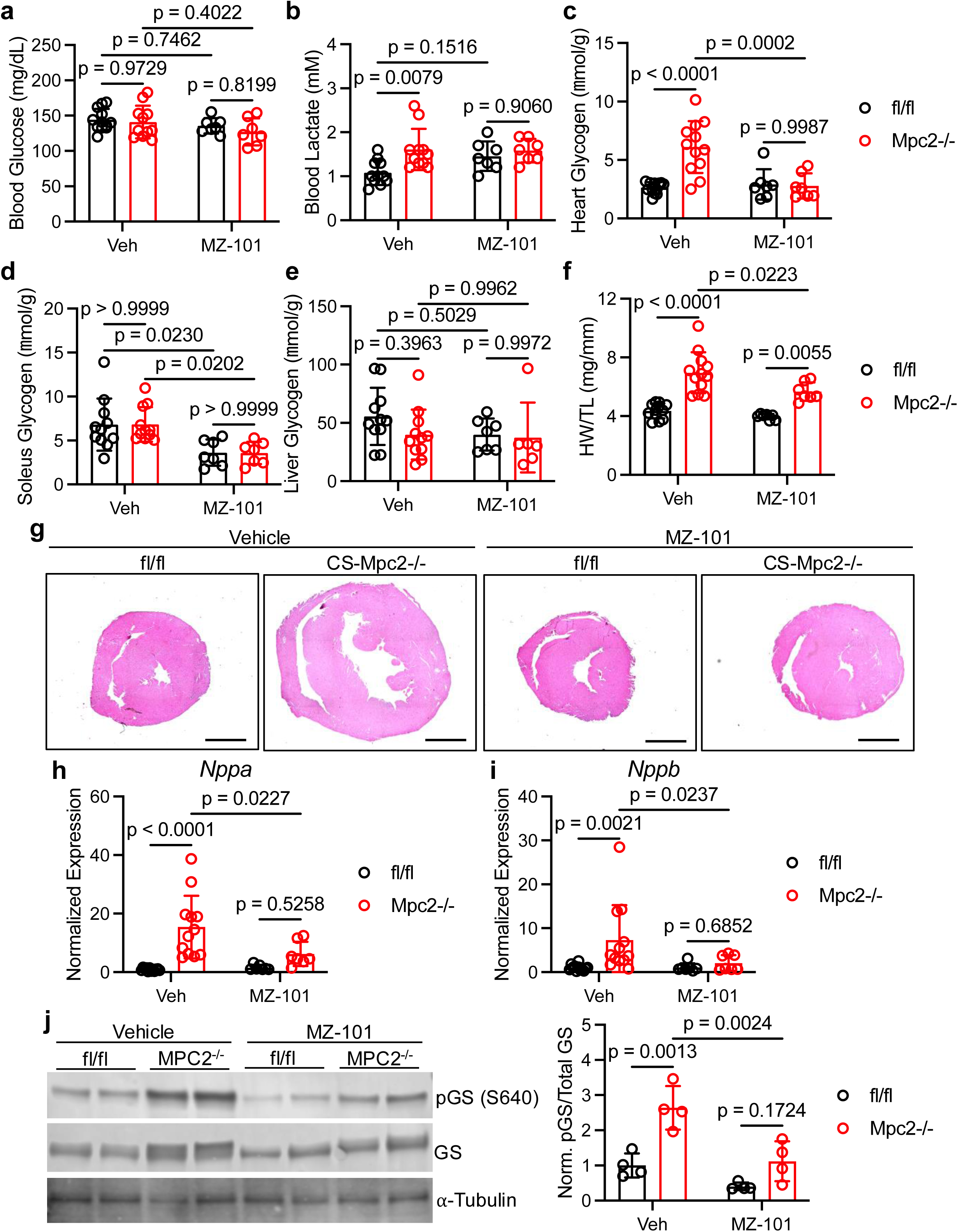
Inhibition of glycogen synthase with MZ-101 reduces glycogen and improves heart failure in CS-Mpc2-/- mice. a-b. Blood glucose **(a)** and blood lactate **(b)** measured after a 2 hour fast from fl/fl and CS-Mpc2-/- mice treated with either vehicle or 100 mg/kg MZ-101 twice daily for 2 weeks, n=7-12. **c-e** Heart **(c)**, soleus **(d)**, and liver **(e)** glycogen content measured biochemically in fl/fl and CS-Mpc2-/- mice treated with either Veh or MZ-101, n=7-12. **f** Heart weight to tibia length ratio (HW/TL) of fl/fl and CS-Mpc2-/- mice treated with either Veh or MZ-101, n=7-12. **g** Representative H&E stained images of mid-ventricular short axis sections of fl/fl and CS-MPC2-/- mice treated with either Veh or MZ-101. Scale bars = 1 mm. **h-i** Cardiac gene expression for *Nppa* and *Nppb* markers of heart failure from fl/fl and CS-Mpc2-/- mice treated with either vehicle or 100 mg/kg MZ-101 twice daily for 2 weeks, n=7-12. **j** Representative western blot images for phosphorylated and total glycogen synthase 1 and α-Tubulin, and normalized quantified densitometry for phosphorylated/total glycogen synthase, n=4 per group. Data are presented as mean±SD, and were analyzed by unpaired, two-tailed Student’s t-test with Welch correction.

### Ketogenic diet normalizes the CS-Mpc2-/- heart failure and glycogen accumulation

Previous reports by us and others showed that feeding a ketogenic diet could either prevent or reverse the heart failure in MPC-deficient hearts^4,13^. Here we again normalized the CS-Mpc2-/- hearts within 3 weeks of ketogenic diet feeding (Fig. 6a). This was associated with complete normalization of the glycogen content of the hearts measured both biochemically and by electron microscopy (Figs. 6b,c). Phosphorylation of glycogen synthase serine-640, which was dramatically increased in the failing CS-Mpc2-/- hearts from low fat (LF)-fed mice, was also normalized by KD-feeding (Fig. 6d). To evaluate glucose metabolism, we once again injected mice with U-^13^C-glucose and measured cardiac metabolite enrichment and pool sizes by mass spectrometry. Interestingly, glucose enrichment was significantly increased in both fl/fl and CS-Mpc2-/- mice fed a KD compared to mice fed LF diet (Fig. 6e), suggesting that these hearts that have been chronically subjected to low blood glucose and insulin from KD feeding^13^ have the ability to enhance glucose uptake when provided with a bolus. This led to increased glycolytic enrichment down to elevated enrichment of pyruvate, lactate, and alanine in hearts from KD-fed mice (Supplementary Figs. 7a-c). However, ^13^C enrichment of TCA cycle intermediates were very low in hearts of KD-fed mice, suggesting that ketogenic diet strongly downregulates glucose carbon incorporation into the TCA cycle and oxidation (Supplementary Figs 7d-l). The failing CS-Mpc2-/- hearts from LF-fed mice once again displayed increased enrichment of the glucose within glycogen, however the increased U-^13^C-glucose uptake with KD-feeding also generally increased ^13^C-enrichment of glycogen (Fig. 6f). However, the metabolite pool sizes of glucose-6-phosphate, pyruvate, and the glycogen precursor UDP-glucose, paint the clear picture of accumulation in the failing CS-Mpc2-/- heart, and normalization with KD-feeding (Figs. 6g-i). Lastly, in confirmation of our biochemical assay and electron micrographs, glucose within the glycogen pool measured by mass spectrometry was significantly elevated in CS-Mpc2-/- hearts from LF-fed mice and normalized by KD-feeding (Fig. 6j). All together, these data suggest that KD rescues the heart failure in CS-Mpc2-/- mice likely by decreasing the need for pyruvate entry into the TCA cycle and normalizing glycolytic metabolism and glycogen storage.

**Fig. 6:**
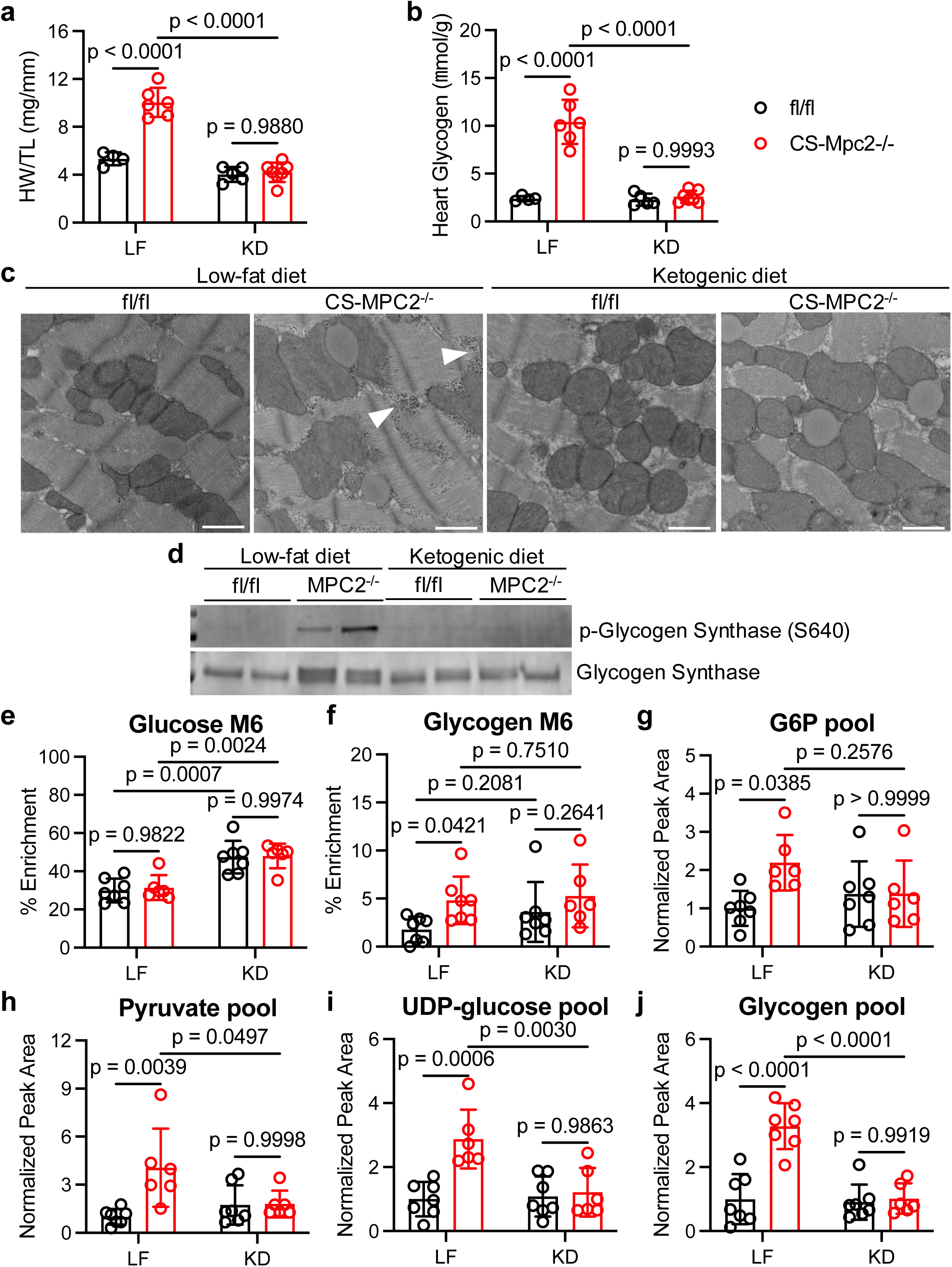
Ketogenic diet normalizes the CS-Mpc2-/- heart failure and glycogen accumulation. **a** Heart weight to tibia length ratio (HW/TL) of fl/fl and CS-Mpc2-/- mice fed either low fat (LF) or ketogenic diet (KD), n=4-7. **b** Heart glycogen content measured biochemically in fl/fl and CS-Mpc2-/- mice fed either a LF diet or KD, n=4-7. **c** Representative transmission electron microscopy images at 5,000X magnification of the cardiac papillary muscle of fl/fl and CS-Mpc2-/- mice fed LF or KD. Arrowheads denote glycogen granules. Scale bars = 1 μm. **d** Western blot images of phosphorylated glycogen synthase (serine 640) and total glycogen synthase from hearts of fl/fl or CS-Mpc2-/- mice fed either LF or KD, n=2. **e-j** ^13^C % enrichment of glucose **(e)** and the glucose within glycogen **(f)**, and the normalized pool size of glucose-6-phosphate (G6P) **(g)**, pyruvate **(h)**, uridine diphosphate (UDP)-glucose **(i)**, and glycogen **(j)** in fl/fl and CS-Mpc2-/- mice that were fed LF or KD and injected i.p. with U-^13^C-glucose, n=6-7. Data are presented as mean±SD, and were analyzed by two-way analysis of variance (ANOVA) with Tukey post-hoc multiple-comparisons test. See also Supplementary Fig. 7.

### AngII-induced cardiac hypertrophy increases glycogen and is improved by ketogenic diet

To evaluate if increased cardiac glycogen was characteristic of non-genetic cardiomyopathy, we subjected mice to AngII infusion via subcutaneous osmotic pump in combination with LF or KD-feeding. As expected, KD-feeding significantly lowered blood glucose concentrations and increased plasma β-hydroxybutyrate levels in both saline and AngII infused mice (Figs. 7a-b). AngII infused mice fed LF displayed significant hypertrophy as shown by increase in HW/TL, which was significantly reduced by KD-feeding (Fig. 7c). Cardiac glycogen more than doubled in AngII infused mice fed LF diet, and again cardiac glycogen levels normalized by KD (Fig. 7d). AngII was previously shown to reduce cardiac MPC1 and MPC2 expression^12^, and we confirmed that both *Mpc1* and *Mpc2* expression were significantly decreased by AngII in LF-fed mice (Figs. 7e-f). Interestingly, ketogenic diet significantly reduced cardiac *Mpc* expression compared to LF (Figs. 7e-f), potentially contributing to the significant decrease of ^13^C-glucose enrichment of the TCA cycle we observed with KD (Supplementary Fig. 7). Like in the failing CS-Mpc2-/- hearts, there was a significant increase in phosphorylation of glycogen synthase in the hearts of AngII infused mice fed LF diet (Figs. 7g-h), suggesting a cellular attempt to inhibit glycogen synthesis. This phosphorylation of glycogen synthase was also slightly increased in hearts from KD-fed mice (Figs. 7g-h), likely due to the low dietary carbohydrate and decreased glycemia (Fig. 7a). All together, these data suggest that AngII-induced cardiac hypertrophy decreases *Mpc* expression and increases glycogen stores; however, this can be normalized through KD-feeding which also improves the cardiac hypertrophy.

**Fig. 7:**
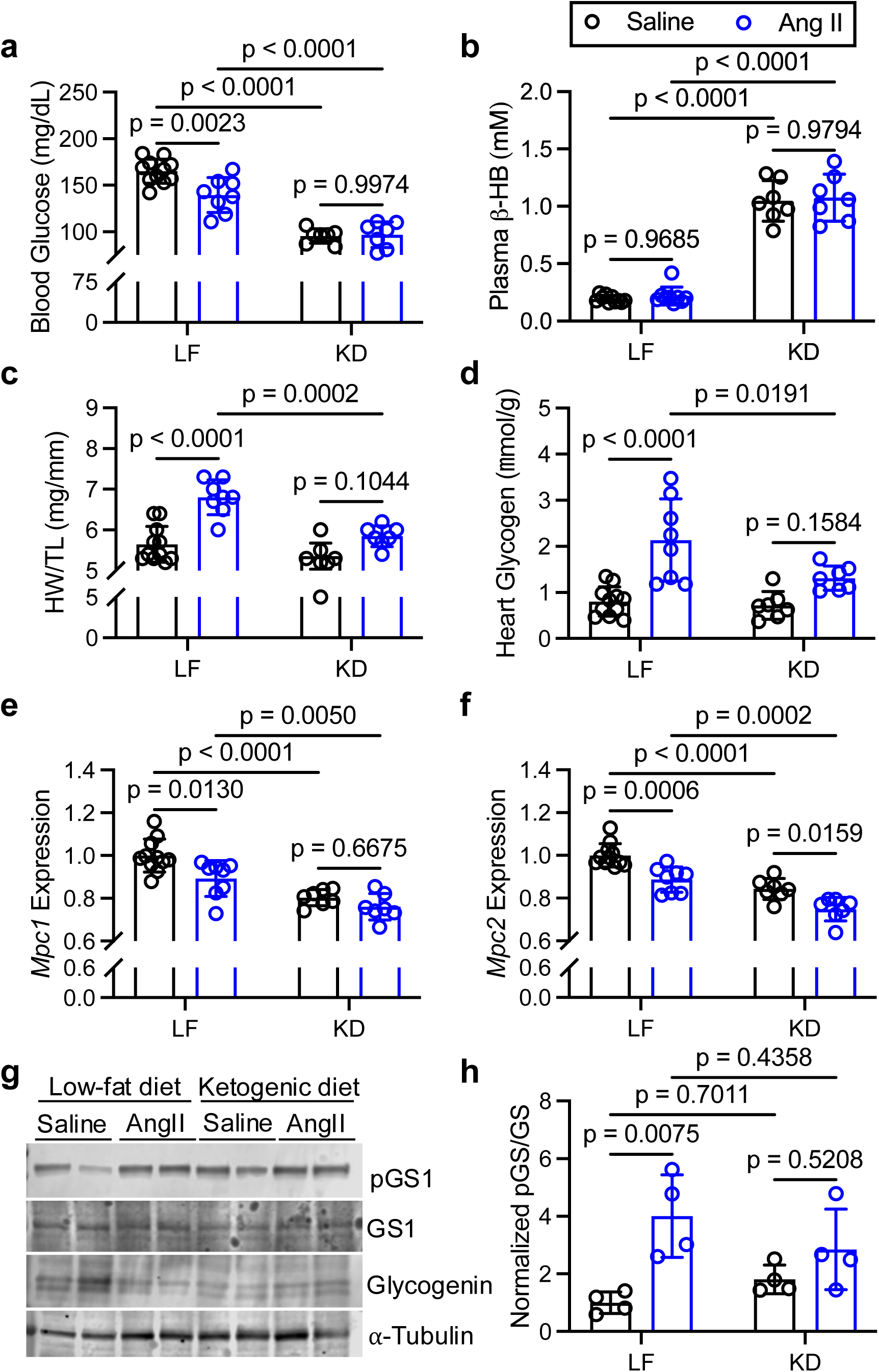
AngII-induced cardiac hypertrophy increases glycogen and is improved by ketogenic diet. **a** and **b** Blood glucose (**a**) and plasma β-hydroxybutyrate (β-HB) concentrations (**b**) of saline vehicle or AngII-infused mice fed low fat (LF) diet or ketogenic diet (KD), n=7-10. **c** Heart weight to tibia length ratio (HW/TL) of saline vehicle and AngII infused mice fed LF or KD, n=7-10. **d** Cardiac glycogen measured biochemically from mice infused with saline or AngII fed either LF or KD, n=7-10. **e** and **f** Normalized gene expression for *Mpc1* (**e**) and *Mpc2* (**f**) in hearts from mice infused with saline or AngII fed LF or KD, n=7-10. **g** and **h** representative western blot images of phosphorylated glycogen synthase (pGS1), total glycogen synthase (GS1), glycogenin-1, and α-tubulin (**g**), and the normalized densitometry quantified for pGS/GS (**h**) from hearts of mice infused with saline or AngII fed LF or KD, n=4. Data are presented as mean±SD, and were analyzed by two-way analysis of variance (ANOVA) with Tukey post-hoc multiple-comparisons test.

## Discussion

This work describes that decreases in MPC expression, or genetic deletion of the MPC, increases cardiac glucose utilization, glycogen accumulation, and heart failure. Altered cardiac metabolism is a hallmark of heart failure, generally consisting of decreased mitochondrial oxidative capacity, particularly for fatty acids which the normal heart prefers^2^. It is generally believed that this decrease in mitochondrial oxidation creates an “energy starved” heart that then causes an increase in myocardial glucose uptake and glycolysis to attempt to fuel the heart^16^. Our current study suggests this may not represent the full story beyond these metabolic changes. In hearts deficient for the mitochondrial pyruvate carrier, which limits the ability of pyruvate from glucose or lactate to enter the mitochondria for oxidation, profound dilated cardiomyopathy develops despite normal levels of high energy phosphates ATP and phosphocreatine. ATP was also previously found to be normal in MPC1-/- hearts that had begun to fail^12^. Although this indicates a lack of “energetic stress” or the “energy starved failing heart”, glucose uptake and glycolysis is increased in these CS-Mpc2-/- hearts. Previous studies suggest that increased glucose utilization is important for fueling the hypertrophied or failing heart, predominantly by increasing nucleotide and amino acid anabolism through the pentose phosphate pathway^7,9^. Here, we suggest that accumulation of glycogen is contributing to the heart failure in CS-Mpc2-/- hearts and in AngII-induced hypertrophy.

The heart has very little capacity to store nutrients or high energy phosphates^24^, making constant tissue perfusion critical. This includes very little storage of glucose as glycogen, with the mouse heart typically containing <5 μmol glucosyl units per gram, compared to 5-30 μmol/g in skeletal muscles and 50-200 μmol/g in the liver in the fed condition^25^. Of course, skeletal muscle glycogen is important during exhaustive exercise, and liver glycogen stores are critical for maintaining glycemia during fasting, and these physiologic stressors can deplete the skeletal muscle and liver glycogen pools. Interestingly, heart glycogen levels remain normal or are even elevated in the fasted state^13,26–28^, with this elevation likely due to the hearts switching to even more predominant fat oxidation in the fasted state inhibiting glycolysis more than glucose uptake^29^. High cardiac workloads increase glycogenolysis and may decrease cardiac glycogen concentrations^30,31^, however these perfused heart studies showed that perfusion with ample substrates (glucose or lactate) abrogated the depletion of glycogen. Thus, it is not well understood if cardiac glycogen levels are decreased or maintained under higher workloads *in vivo*. Exhaustive exercise has been shown to decrease cardiac glycogen content by 25%, which was rapidly repleted^32^. Overall, these previous studies suggest that while cardiac glycogen levels are low, the consistent flux of glucose in and out of the glycogen pool and relative maintenance of glycogen concentrations indicates an important role.

Another interesting point from these previous studies is that glucose released from cardiac glycogen is preferentially oxidized instead of converted to lactate^30,33^. This would have significant effects in MPC-deficient hearts that cannot efficiently oxidize this glycogen-derived pyruvate. This inability to oxidize glucose from glycogen may partially explain the accumulation of glycogen in these MPC-deficient hearts, in addition to the increased glucose uptake and incorporation of ^13^C-glucose into glycogen we observed. That said, a limitation of our current study was the inability to assess rates of glycogenolysis.

A puzzling finding from this study, and a previous study^4^, is that enrichment of the TCA cycle from ^13^C-glucose is not decreased in MPC-/- hearts (Supplementary Figs. 2 and 6), even though we observe significantly reduced respiration with pyruvate as substrate in isolated mitochondria and permeabilized cardiac muscle fibers from CS-Mpc2-/- hearts (Supplementary Fig. 3). There are several potential explanations for this maintained TCA cycle enrichment from ^13^C-glucose. In this whole-heart method, ^13^C enrichment of the TCA cycle would be maintained in the non-cardiomyocyte cells in the heart which would not have *Mpc2* deleted by Mlc2vCre. This explanation is supported in that decreased TCA cycle enrichment was observed in isolated cardiomyocytes from MPC1-/- hearts^14^. Another possibility is that there could be a metabolic bypass of the MPC, such as alanine transamination and alanine mitochondrial transport as we have previously identified in hepatocytes^17^, or mitochondrial import of lactate as recently suggested^18^. This sort of bypass may be occurring to a degree, as we do observe much greater reduction of pyruvate respiration in isolated mitochondria (devoid of cytosolic enzymes) than we do when using permeabilized cardiac tissue (Supplementary Fig. 3). Lastly, while we mostly think of these organic acids as TCA cycle metabolites, it is clear many of these can be released from mitochondria and found in the cytosol and other organelles^34^. Therefore, non-mitochondrial pools of these metabolites may somehow be labeled by ^13^C-glucose in the CS-Mpc2-/- hearts. Overall, with the dramatic reduction in mitochondrial pyruvate respiration and the pronounced pathologic phenotype of the CS-Mpc2-/- hearts, we infer that pyruvate oxidation is reduced despite normal ^13^C enrichment of TCA cycle metabolites in these studies.

Perhaps due to its low abundance, cardiac glycogen has not been well studied in the context of heart failure. The one exception is the association between glycogen accumulation and hypertrophic cardiomyopathy that occurs in multiple forms of glycogen storage diseases (GSD). For example, GSD type-II, or Pompe disease, is caused by mutation in the enzyme acid _a_-glucosdiase, which catalyzes the lysosomal degradation of glycogen^35^. This results in cardiac glycogen accumulation, up to 200-fold in Pompe disease mouse models, and development of hypertrophic cardiomyopathy^36^. Currently, Pompe disease treatment involves enzyme replacement therapy, requiring regular invasive and time-consuming infusions. Recent studies evaluated MZ-101, a glycogen synthase 1 inhibitor, in mouse models of Pompe disease which significantly reduced glycogen load in skeletal and cardiac muscle^22,23^. Other than reduction in glycogen, cardiac endpoints were not evaluated in either study. These two studies suggest the potential efficacy of “substrate reduction therapies” for Pompe disease^22,23^. Interestingly, both these mouse models of Pompe disease^22^ and the CS-Mpc2-/- hearts in which glycogen accumulates display increased phosphorylation of glycogen synthase which should be inhibitory, yet glycogen synthase activity remains or is enhanced in these pathologies. Thankfully, MZ-101 inhibits both the unphosphorylated and phosphorylated forms of GYS1^22^. In summary, glycogen storage diseases clearly support the connection between increased cardiac glycogen and heart failure, and this current work suggests that cardiac glycogen reduction with MZ-101 successfully improves heart failure caused by defective mitochondrial pyruvate metabolism.

This current work also describes the normalization of cardiac glycogen and rescuing of heart failure by a KD in both the CS-Mpc2-/- model, and in AngII-induced hypertrophy. We hypothesize that KD is improving these hearts by decreasing glycemia and cardiac glucose metabolism, ultimately reducing the need for mitochondrial pyruvate metabolism and the MPC as shown by very low ^13^C-glucose enrichment of the TCA cycle in hearts from KD-fed mice (Supplementary Fig. 7). Significantly reduced glucose oxidation, even with insulin stimulation, was recently observed in hearts from mice fed a KD^37^. One interesting point to this current study however is that hearts from KD-fed mice appeared to have increased uptake of the bolus ^13^C-glucose (Fig. 6e). This is somewhat contrary to the KD-induced insulin resistance recently described^37^, and argues that the hearts of KD-fed mice can readily utilize glucose when challenged with a bolus. Ultimately, this chronic decrease in glycemia and cardiac glucose metabolism from ketogenic diet feeding allows the glycogen levels to return to normal. Thus, our findings not only provide new insights into how reduced mitochondrial pyruvate metabolism contributes to heart failure but also provides important implications for the development of strategies to reduce cardiac glucose utilization and glycogen levels. One could speculate this decrease in glycogen may also be a mechanism for the profound cardioprotective effects of sodium-glucose cotransporter-2 (SGLT2) inhibitors, as recently postulated that SGLT2 inhibitors improve heart failure by reducing intermediates of glucose and lipid metabolism^38^. However, cardiac glycogen content with SGLT2 inhibitor treatment requires further investigation.

## Methods

### Animals

All animal procedures were performed in accordance with National Institutes of Health guidelines and approved by the Institutional Animal Care and Use Committee at Saint Louis University. Generation of mice with *Mpc2* gene flanked by loxP sites, as well as cardiac-specific deletion of *Mpc2* by crossing to Mlc2v-Cre knock-in mice (Jackson Laboratory Strain #029465) have been described previously^13,17^. All mice used in these experiments were from a C57BL6/J background. All experiments were performed with a mixture of male and female littermate mice. Mice were housed in a climate-controlled specific-pathogen free facility maintained at 22-24°C and 40-60% humidity in ventilated cages with a 12-hour light/dark cycle. Ad libitum access to drinking water was provided by automatic cage waterers. Food was also provided ad libitum. All mice were group-housed, up to five mice per cage. Based on our previous study^13^, the experiments in young, non-failing CS-Mpc2-/- mice were performed at 6 weeks of age and experiments in CS-Mpc2-/- mice with heart failure were performed at 14-30 weeks of age. For ketogenic diet (KD) experiments, fl/fl and CS-Mpc2-/- mice were initially fed chow and aged to 16 weeks to develop heart failure in CS-Mpc2-/- mice, then switched to either low-fat (LF) control diet or KD for a duration of 3 weeks. The LF diet (F1515, BioServ) composition is 68.9%kcal carbohydrate, 19.0%kcal protein, and 12.1%kcal fat. Ketogenic diet (F3666, BioServ) composition is 1.8%kcal carbohydrate, 4.7%kcal protein, and 93.5%kcal fat. Diets were randomly assigned for whole cages based on genotypes contained within to result in roughly even numbers for each group. Unless otherwise stated, at study culmination mice were fasted for 2-3 hours, and euthanized by CO_2_ asphyxiation. Blood was collected via cannulation of the inferior vena cava and placed into EDTA-treated tubes on ice. Tissues were then excised, weighed, and snap frozen in liquid nitrogen. Plasma was collected by centrifuging blood tubes at 8,000*g* for 8 minutes at 4°C and freezing the plasma supernatant in liquid nitrogen. Plasma and tissues were stored at -80°C until analyzed.

MZ-101 was purchased from Enamine US, Inc, and was dissolved in 45% H2O, 30% polyethylene glycol 200, 20% polypropylene glycol, and 5% N-methyl-2-pyrrolidone (all from Sigma Aldrich) as performed previously ^22^. Gavage dosing (100 mg/kg MZ-101 or similar volume of vehicle) was performed twice daily in 14-week-old CS-Mpc2-/- or fl/fl littermate mice. Fresh gavage solutions were prepared weekly. Mice were treated for 2 weeks, then euthanized by CO_2_ and plasma/tissues harvested as described above.

AngII infusion was performed in 9-week-old wildtype C57BL6/J male mice purchased from the Jackson Laboratory (Strain #000664). Subcutaneous osmotic pumps (#1004, Alzet) with either AngII (1.44mg/kg/day, Sigma Aldrich) or saline vehicle were implanted for an infusion duration of three weeks similarly to as described previously^39^. Mice were anesthetized by inhaled 2-3% isoflurane in 100% oxygen via a nose cone, and the upper back area shaved with electric clippers and scrubbed with 7.5% betadine antiseptic. A ∼1cm incision was made in the prepped area of the mouse’s back, and forceps inserted to create a subcutaneous pocket. After inserting the pump, the wound was sealed with 2-3 surgical staples. Mice were treated with 1 mg/kg subcutaneous buprenorphine-SR and several drops of lidocaine were applied to wound edges for pain management. Mice were allowed to arouse while on a heat pad and placed back into their home cage. Mice were observed daily, and surgical staples were removed within 7-10 days. Mice were switched to LF or KD 3 days prior to pump implant. Diet was assigned randomly for each cage, and then vehicle versus AngII pumps were assigned for mice for each diet to result in roughly even numbers per group. Mice were euthanized by CO_2_ and plasma/tissues harvested as described above.

### RNA sequencing and analysis

Hearts used for RNA sequencing were excised from 6 fl/fl and 6 CS-Mpc2-/- 30-week-old chow-fed mice. RNA was isolated from ∼25 mg of frozen left ventricular tissue using RNA-STAT (Tel-Test, Inc) and purified using the PureLink RNA Mini Kit with DNase treatment (Invitrogen). RNA was then pooled between 2 different fl/fl and CS-Mpc2-/- hearts, resulting in n=3 samples per group. RNA concentration and quality were determined by 2100 Bioanalyzer RNA 6000 Nano Kit (Agilent) and depleted of rRNA using RiboMinus Eukaryote Kit v2 (Ambion). DNA libraries were constructed using Ion Total RNA-seq Kit v2 (Thermo Fisher Scientific) and sequenced on an Ion Torrent Proton DNA sequencer (Thermo Fisher Scientific). Raw data were aligned to the mm10 genome. Gene counts for each sample were normalized by read depth, low expression genes filtered from the data, and *t*-test performed for statistical comparison. Lists of upregulated and downregulated differentially expressed genes (p < 0.05) between fl/fl and CS-Mpc2-/- hearts were subjected to pathway-level overrepresentation analysis using IMPaLA^40^, which included Kyoto encyclopedia of genes and genomes (KEGG) pathway analysis, among others. Heat map of differentially expressed genes was generated with Shiny heatmap^41^. RNA sequencing files were uploaded to NCBI GEO Series Number GSE268694.

### Gene expression analysis

Relative gene expression was determined by reverse transcription quantitative PCR (RT-qPCR). Total RNA was extracted from snap frozen tissues using RNA-STAT (Tel-Test, Inc). [RNA] was assessed by Nanodrop 2000c (Thermo Fisher Scientific). Then 1 _µ_g of sample was reverse transcribed into complementary DNA by Superscript VILO (Thermo Fisher Scientific) using a 2720 Thermal cycler (Applied Biosystems). Relative quantification of target gene expression was measured in duplicate using Power SYBR Green (Thermo Fisher Scientific) with a QuantStudio 3 PCR System (Thermo Fisher Scientific). Target gene threshold cycle (Ct) values were normalized to reference gene (*Rplp0*) Ct values by the 2^-ΔΔCt^ method. Oligonucleotide primer sequences used in 5’ to 3’ were: *Rplp0* FWD: GCAGACAACGTGGGCTCCAAGCGAA, *Rplp0* REV: GGTCCTCCTTGGTGAACACGAAGCCC, *Mpc1* FWD: GACTTTCGCCCTCTGTTGCTA, *Mpc1* REV: GAGGTTGTACCTTGTAGGCAAAT, *Mpc2* FWD: CCGCCGCGATGGCAGCTG, *Mpc2* REV: GCTAGTCCAGCACACACCAATCC, *Nppa* FWD: AGTGCGGTGTCCAACACAGA, *Nppa* REV: GACCTCATCTTCTACCGGCATCT, *Nppb* FWD: GCTGCTTTGGGCACAAGATAG, and *Nppb* REV: GCAGCCAGGAGGTCTTCCTA.

### Western blotting and protein expression analysis

Protein extracts were prepared by homogenizing ∼30 mg of frozen ventricular tissue in lysis buffer (15mM NaCl, 25 nM Tri Base, 1 mM EDTA, 0.2% NP-40, 10% glycerol) supplemented with 1X cOmplete protease inhibitor cocktail (Millipore Sigma) and a phosphatase inhibitor cocktail (1 mM Na_3_VO_4_, 1 mM NaF, and 1 mM PMSF) by using 2-3 zirconia beads (Biospec products) and homogenization for 1 min with a Mini Beadbeater (Biospec products). Protein concentrations were measured using MicroBCA Assay kit (Thermo Fisher Scientific) and absorbance measured with a Synergy HTX multi-mode plate reader (Biotek). Then 50 μg of protein lysate was electrophoresed on criterion TGX 4-15% precast gel (Bio-Rad Laboratories) and transferred onto PVDF membrane (Millipore). Membranes were blocked with 5% bovine serum albumin (BSA) (#BAH65 Equitech-Bio, Inc) in Tris-buffered saline with Tween-20 (TBST) for at least 1 hour.

Primary antibodies were used at 1:1000 dilution in 5% BSA-TBST, with gentle rocking at 4°C overnight. AKT (9272), phosphorylated AKT (serine 473) (4060), MPC1 (14462), MPC2 (46141), glycogen synthase (3886), and phosphorylated glycogen synthase (serine 640) (47043) were from Cell Signaling. α-tubulin (T6199) was from Sigma. PYGM (ab231963) was from abcam. Glycogenin-1 (sc-271109) was from Santa Cruz Biotechnology. After primary antibody incubation, membranes were washed 3X for 10 minutes in TBST and probed with IRDye secondary antibodies at 1:10,000 (Li-Cor Biosciences, 926-32213 or 926-68072) in 5% BSA-TBST for 1 hour, then washed 3X 10 min. Membranes were imaged on an Odyssey CLx imaging system and analyzed with Image Studio Lite software (Li-Cor Biosciences).

### Targeted metabolomics and ^13^C enrichment from glucose

Mice were fasted for 2 hours then injected intraperitoneally (i.p.) with 1g/kg uniformly labelled U-^13^C-glucose (CLM-1396, Cambridge Isotope Labs) in saline. 30 min later the mice were anesthetized with 1-2% isoflurane (Covetrus) in 100% oxygen via an induction chamber then transferred to a nose cone. A thoracotomy was rapidly performed and the beating heart snap-frozen using a metal clamp chilled in liquid nitrogen. Hearts were stored at -80°C until processed. The heart tissue was pulverized with a mortar and pestle chilled in a liquid nitrogen bath. To every 1 mg of pulverized tissue, 40 μl of 2:2:1 acetonitrile:methanol:water was added. The samples were then briefly vortexed, placed in a liquid nitrogen bath for 1 minute, and then sonicated for 10 minutes at 25°C. This was repeated 3X. Samples were stored at -20°C overnight. The following day, the samples were centrifuged at 14,000*g* for 10 min at 4°C, and supernatants added to glass liquid chromatography vials with autosampler inserts and stored at -80°C until analyzed.

Ultra-high-performance liquid chromatography (UHPLC)-MS/MS was performed with a Vanquish Flex UHPLC system interfaced with an Orbitrap ID-X Tribid Mass spectrometer (Thermo Fisher Scientific) similar to as previously described^42^. Hydrophilic interaction liquid chromatography (HILIC) separation was performed using a HILICON iHILIC-(P) Classic column (Tvistevagen, Umea, Sweden) with the following specifications: 100 × 2.1 mm, 5 μm. Mobile-phase solvents were composed of A = 20 mM ammonium bicarbonate, 0.1% ammonium hydroxide and 2.5 μM medronic acid in water:acetonitrile (95:5) and B = acetonitrile:water (95:5). The column compartment was 45°C for all experiments. The following linear gradient was applied at a flow rate of 250 μL min ^−1^: 0–1 min: 90% B, 1–12 min: 90-35% B, 12–12.5 min: 35-25% B, 12.5–14.5 min: 25% B. The column was re-equilibrated with 20 column volumes of 90% B. The injection volume was 4 μL for all experiments.

Data were collected with the following settings: spray voltage, 3.5kC (positive) / −2.8 kV (negative); sheath gas, 50; auxiliary gas, 10; sweep gas, 1; ion transfer tube temperature, 300°C; vaporizer temperature, 200°C; mass range, 67–1,000 Da, resolution, 120,000 (MS1), 30,000 (MS/MS); maximum injection time, 100 ms; isolation window, 1.6 Da. LC/MS data were processed and analyzed with the open-source Skyline software^43^. Natural abundance correction of ^13^C for tracer experiments was performed with AccuCor^44^. All studies contained 3 quality control samples and a blank sample, and only metabolites with sufficient abundance and fractional enrichment over the limit of detection were included in analysis. These methods were used to analyze the pool size and ^13^C enrichment of glucose and glycolytic metabolites, organic acids and amino acids, and the pool sizes of nucleotide phosphates and phosphocreatine and creatine. Data are presented as the % enrichment of the specified ^13^C isotopologue, or as the normalized peak area for each metabolite’s pool size.

### Transmission electron microscopy and gross histology

The papillary muscle from hearts (under 1 mm^3^) was excised and fixed in 2.5% glutaraldehyde in 0.1M sodium cacodylate, pH 7.4 overnight. Samples were then post-fixed with 1% osmium tetroxide for 1 h and then serial dehydration with alcohol (50%, 75%, 90%, and 100%) was performed and then embedded in Epon 12. Ultrathin sections were then post-stained with uranyl acetate and lead citrate. Samples were imaged on a JEOL 1400 Plus transmission electron microscope equipped with an AMT camera.

Hearts from vehicle and MZ-101 treated mice were cut into a ∼1 mm short axis slice with a razor blade and fixed in 10% neutral buffered formalin. Slices were then processed, embedded, sectioned, and stained by H&E by the Saint Louis University Advanced Spatial Biology and Research Histology Core.

### Cardiac glycogen assay

Cardiac glycogen concentrations were measured similarly to as preformed previously^13^. Briefly, 15-30 mg of heart tissue were boiled in 30% KOH at 100°C for 30 min. Then, 2% Na_2_SO_4_ and 100% EtOH were added, and the tubes were vortexed. Samples were boiled again to help aid in the precipitation of glycogen and then tubes were centrifuged at 16,000*g* for 5 min and supernatant aspirated. The pellet was dissolved in 200 μL H_2_O and 800 μL of 100% ethanol was added. The samples were then boiled for 5 minutes, and recentrifuged at 16,000*g* for 5 min. This wash step was repeated 3X. The final pellets were suspended in 200 μL of 0.3 mg/ml amyloglucosidase (Roche) in 0.2 M sodium acetate (pH 4.8). Serial dilutions of 10 mg/ml oyster glycogen (Sigma) were prepared as standards. Samples and standards were then incubated in a 40°C water bath for 3 hours. Samples and standards were then diluted 1:1 with H_2_O, and the samples were added to a 96-well plate. Glucose assay buffer (0.3 M triethanolamine, 4 mM MgCl_2_, 2 mM ATP, 0.9 mM NADP^+^, and 5 μg/mL glucose-6-phosphate dehydrogenase (Roche)) was added to each well, and the absorbance was measured at 340 nm with a Synergy HTX multi-mode plate reader (Biotek). Then 1 μg hexokinase (Roche) was added to each well and the plate was incubated at room temperature in the dark for 30 minutes, and absorbance measured again at 340 nm. The glycogen concentration was calculated from the difference in absorbance readings plotted in relation to the oyster glycogen standards.

Glycogen ^13^C enrichment and pool size measurement was performed similarly as described previously^22^. Briefly, glycogen from cardiac tissue of mice injected with U-^13^C-glucose as described above was precipitated and digested by 0.3 mg/mL amyloglucosidase in 0.2 M sodium acetate as described for the biochemical glycogen assay. Following the 3 hour digestion with amyloglucosidase, the samples were added to a 2:2:1 acetonitrile:methanol:water mixture maintaining a 1:9 ratio of sample: extraction solvent. The samples were vortexed for 1 min then stored at -20°C for 1 hour. Then the samples were centrifuged at 14,000*g* for 10 min at 4°C. The supernatant was then transferred to glass LC vials for UHPLC-MS/MS analysis as described above.

### Myocardial glucose uptake assay

Glucose uptake was measured similarly to methods used for skeletal muscle and papillary muscle as described previously^45–47^. Mice were anesthetized with 50 mg/kg sodium pentobarbital intraperitoneal (i.p.) injection, the heart removed, and two ∼2-4 mg left ventricular muscle strips collected from each heart. Individual cardiac strips were placed into a well of a 24-well plate containing Krebs-Henseleit Buffer recovery solution with non-glucose fuel sources (4.5 mM lactate, 0.5 mM pyruvate, and 200 μM BSA-palmitate), 34.8 mM mannitol, and 0.1% radioimmunoassay-grade BSA at 35°C. Incubation buffers were pre-gassed with 95% O_2_:5% CO_2_, and each well of the 24-well plate had its own gas supply. After 60 min in recovery buffer, samples were moved to another well containing the same solution ± 30 μU/mL (submaximal) insulin for 30 min. Samples were then moved to another well containing glucose transport medium ± 30 μU/mL insulin. Glucose transport assay medium was the same as recovery medium but also contained 1 mM 2-deoxyglucose (2-DG), 2 μCi/mL ^3^H-2-DG, and 0.3 mCi/mL ^14^C-mannitol at 30°C. After 10 min in the glucose transport assay medium, samples were clamp frozen. Samples were homogenized in ice-cold buffer containing protease and phosphatase inhibitors (50 mM HEPES, pH 7.4, 2 mM Na_3_VO_4_, 150 mM NaF,10 μg/mL leupeptin, 10 μg/mL aprotinin, 0.5 μg/mL pepstatin and 1 mM phenylmethylsulfonyl fluoride) and centrifuged at 4°C for 10 min at 13,500*g*. Supernatant protein was assessed by the Bradford method, and aliquots were mixed with scintillation fluid (Ultima Gold XR, Perkin Elmer, Boston, MA, USA) before counting of disintegrations per minute (DPM, TriCarb 3110TR, Perkin Elmer). The DPM of ^14^C mannitol was used to measure extracellular volume and correct for DPM of ^3^H-2-DG in the extracellular fluid.

### Insulin signaling assessment

Insulin signaling was evaluated as described previously^48^. Briefly, mice were fasted for 3 hours, then injected i.p. with 5mU/g insulin or saline vehicle. 10 minutes after injection, mice were euthanized by CO_2_ and the hearts collected and snap frozen in liquid nitrogen and stored at -80°C. Western blotting was then performed for phosphorylated and total AKT as described above.

### Plasma ketone body assay

Plasma ketone concentrations were measured using a β-hydroxybutyrate colorimetric assay kit (Cayman Chemical). Samples were incubated in the dark for 25 min and absorbance measured at 445 nm with a Synergy HTX multi-mode plate reader (Biotek) and plotted against a standard curve for quantification.

### Mitochondrial isolation and respirometry

Mitochondria were isolated from freshly excised hearts by ten passes of a glass-on-glass Dounce homogenizer on ice in 4 mL of buffer containing 250 mM sucrose, 10 mM Tris, and 1 mM EDTA, pH 7.4. Homogenates were centrifuged at 1,000*g* x 5 min at 4°C, and the supernatant transferred to a new tube and centrifuged at 10,000*g* x 10 min at 4°C. This mitochondria-enriched pellet was washed with fresh homogenization buffer and recentrifuged at 10,000*g* x 10 min at 4°C twice. The mitochondrial pellet was then solubilized in ∼200 μL of Mir05 respiration buffer consisting of 0.5 mM EGTA, 3 mM MgCl_2_, 60 mM lactobionic acid, 20 mM taurine, 10 mM KH_2_PO_4_, 20 mM HEPES, 110 mM sucrose, and 1g/L fatty-acid free BSA, pH 7.1. Mitochondrial protein concentration was measured by MicroBCA (Thermo Fisher Scientific).

50 μg of mitochondria were added to each chamber of an Oxygraph O2k FluoRespirometer (Oroboros Instruments), and respiration was stimulated with 5 mM pyruvate, 2 mM malate, and then coupled with 2.5 mM ADP+Mg^2+^. Lastly, 5 μM UK-5099 was added to the chambers to inhibit the MPC and pyruvate respiration.

Respiration was also performed on saponin-permeabilized left-ventricular muscle fibers using the Oroboros O2k similar to as previously performed^13^. Respiration was again stimulated with 5 mM pyruvate, 2 mM malate, and then coupled with 2.5 mM ADP+Mg^2+^, and inhibited with 5 μM UK-5099. Respiration data was recorded and analyzed with DatLab version 7.4.0.1 software, and is presented as oxygen consumption rate normalized to mitochondrial mass or tissue mass added to the chamber.

### Statistical analysis

All data are presented as dot plots with mean±S.D. All data were graphed and statistically analyzed with GraphPad Prism version 10.4.2. An unpaired, two-tailed Student’s *t*-test with Welch correction was used for comparison of two groups. Experiments with more than two groups were analyzed using a two-way analysis of variance (ANOVA) with Tukey post-hoc multiple-comparisons test. A *P* value of less than 0.05 was considered statistically significant.

## Nonstandard Abbreviations and Acronyms

2-DG: 2-deoxyglucose
αKG: alpha-ketoglutarate
ADP: adenosine diphosphate
AMP: adenosine monophosphate
ATP: adenosine triphosphate
AngII: angiotensin-II
Cr: creatine
DHAP: dihydroxyacetone phosphate
FAO: fatty acid oxidation
FBP: fructose bisphosphate
G6P: glucose-6-phosphate
GSD: glycogen storage disease
KD: ketogenic diet
KEGG: Kyoto encyclopedia of genes and genomes
LF: low-fat
MPC: mitochondrial pyruvate carrier
NAD+ and NADH: oxidized and reduced nicotinamide adenine dinucleotide
NADP+ and NADPH: oxidized and reduced nicotinamide adenine dinucleotide phosphate
PCr: phosphocreatine
PEP: phosphoenolpyruvate
P/M: pyruvate/malate
R5P: ribose-5-phosphate
RT-qPCR: reverse transcription quantitative PCR, Sedo7P, sedoheptulose 7-phosphate
UDP: uridine diphosphate
UHPLC: ultra-high performance liquid chromatography

## Data availability

Data supporting the findings of this study are available from the corresponding author upon reasonable request. The RNA sequencing data generated in this study have been deposited in NCBI GEO Series Number GSE268694.

## Acknowledgements

R.C.W. is supported by a Doisy Scholar Fellowship from the Doisy Fund of the Edward A. Doisy Department of Biochemistry and Molecular Biology at Saint Louis University School of Medicine. This work was supported by NIH R00-HL136658 to K.S.M and R35 ES028365 to G.J.P.

## Disclosures

G.J.P. has a research collaboration agreement with Thermo Fisher Scientific and is a scientific advisor for Cambridge Isotope Laboratories. K.S.M. receives research support from Cirius Therapeutics and serves as a consultant for BioGenerator Ventures. All other authors declare no conflicts of interest.

## Author Information

Authors and Affiliations

**Edward A. Doisy Department of Biochemistry and Molecular Biology, Saint Louis University School of Medicine, St. Louis, MO, USA**

Rachel Weiss, Kelly Pyles, Michelle Brennan, & Kyle McCommis

**Department of Biology, Saint Louis University, St. Louis, MO, USA**

Jonathan Fisher

**Departments of Chemistry, Medicine, and the Center for Mass Spectrometry and Metabolic Tracing, Washington University in St. Louis, St. Louis, MO, USA**

Kevin Cho & Gary Patti

## Contributions

R.C.W, J.S.F, G.J.P., and K.S.M. conceived and designed the experiments. R.C.W., K.D.P., K.C., M.B., and J.S.F. performed the experiments and analyzed the data. R.C.W and K.S.M. wrote the manuscript. All authors read, edited, and approved the manuscript.

## Figure Legends

**Supplementary Fig. 1: Glycolytic pool sizes in failing MPC-/- hearts.** Related to Fig. 2. **a** Schematic pathway of glycolysis and accessory glucose pathways. **b-l** Normalized pool size of glucose **(b)**, glucose-6-phosphate (G6P) **(c)**, fructose bisphosphate (FBP) **(d)**, dihydroxyacetone phosphate (DHAP) **(e)**, 2/3-phosphoglycerate (PG) **(f)**, ^13^ phosphoenolpyruvate (PEP) **(g)**, pyruvate **(h)**, lactate **(i)**, alanine **(j)**, and the ^13^C % enrichment of glycerol 3-phosphate (Glycerol 3P) (**k**) and glycerol 3P pool **(l)** of failing CS-Mpc2-/- hearts and fl/fl littermates injected with U-^13^C-glucose, n=11-12. Data are presented as mean±SD. Data were evaluated by unpaired, two-tailed Student’s t-test with Welch correction.

**Supplementary Fig. 2: TCA cycle enrichment and pool sizes in failing CS-Mpc2-/-hearts.** Related to Fig. 2. **a-l** the ^13^C % enrichment and pool sizes of citrate **(a-b)**, aconitate **(c-d)**, α-ketoglutarate (KG) **(e-f)**, succinate **(g-h)**, fumarate **(i-j)**, and malate **(k-l)** in fl/fl and failing CS-Mpc2-/- hearts of mice injected i.p. with U-^13^C-glucose, n=11-12. Data are presented as mean±SD. Data were evaluated by unpaired, two-tailed Student’s t-test with Welch correction.

**Supplementary Fig. 3: MPC deletion reduces pyruvate oxidation.** Related to Fig. 2. **a-b** Oxygen consumption rates (OCR) measured from isolated cardiac mitochondria (**a**) and permeabilized cardiac muscle fibers (**b**) from hearts of fl/fl and CS-Mpc2-/- littermates stimulated with pyruvate/malate (P/M), P/M plus adenosine diphosphate (ADP), and P/M, ADP, and the MPC inhibitor UK-5099 (5 μM), n=4-5. Data are presented as mean±SD. Data were evaluated by unpaired, two-tailed Student’s t-test with Welch correction.

**Supplementary Fig. 4: No major changes in the pentose phosphate pathway in failing MPC-/- hearts.** Related to Fig. 2. **a** Schematic of ^13^C enrichment of the pentose phosphate pathway from U-C-glucose. Metabolites in grey were not measured. **b-f** ^13^C % enrichment and normalized pool size of 6-phosphogluconate (PG) **(b)**, ribose-5-phosphate (R5P) **(c)**, sedoheptulose 7-phosphate (Sedo7P) **(d)**, serine **(e)**, and glycine **(f)**, in hearts from fl/fl and failing CS-Mpc2-/- mice injected with U-^13^C-glucose, n=11-12. Data are presented as mean±SD. Data were evaluated by unpaired, two-tailed Student’s t-test with Welch correction.

**Supplementary Fig. 5: Glycolytic pool sizes in young, non-failing MPC-/- hearts.** Related to Fig. 4. **a** Schematic of glycolysis and accessory glucose pathways. **b-q** Normalized pool sizes of glucose **(b)**, glucose-6-phosphate (G6P) (**c**), fructose bisphosphate (FBP) (**d**), dihydroxyacetone phosphate (DHAP) **(e)**, 2/3 phosphoglycerate (PG) **(f)**, phosphoenolpyruvate (PEP) **(g)**, pyruvate **(h)**, lactate **(i**), alanine (j), ribose-5-phosphate (R5P) **(k)**, and the ^13^C enrichment and pool size of serine **(l-m)**, glycine **(n-o)**, and glycerol 3-phosphate (glycerol 3P) **(p-q)** in young non-failing CS-Mpc2-/- and fl/fl littermates injected with U-^13^C-glucose, n=8-10. Data are presented as mean±SD. Data were evaluated by unpaired, two-tailed Student’s t-test with Welch correction.

**Supplementary Fig. 6: TCA cycle enrichment and pool sizes in young, non-failing CS-Mpc2-/- hearts.** Related to Fig. 4. **a-l**^13^C % enrichment and normalized pool sizes of citrate (**a-b**), aconitate (**c-d**), α-ketoglutarate (αKG) (**e-f**), succinate (**g-h**), fumarate (**i-j**), and malate (**k-l**), in hearts of young, nonfailing CS-Mpc2-/- and fl/fl littermates injected with U-^13^C-glucose, n=8-10. Data are presented as mean±SD. Data were evaluated by unpaired, two-tailed Student’s t-test with Welch correction.

**Supplementary Fig. 7: Ketogenic diet decreases TCA cycle enrichment from glucose.** Related to Fig. 6. **a-l** ^13^C % enrichment of pyruvate (**a**), lactate (**b**), alanine (**c**), citrate (**d**), aconitate (**e**), α-ketoglutarate (αKG) (**f**), glutamate (**g**), glutamine (**h**), succinate (**i**), fumarate (**j**), malate (**k**), and aspartate (**l**) in hearts from fl/fl and CS-Mpc2-/- fed either low-fat (LF) or ketogenic diet (KD) and injected with U-^13^C-glucose, n=6-7. Data are presented as mean±SD. Data were evaluated by two-way analysis of variance (ANOVA) with Tukey post-hoc multiple-comparisons test.

## Notes

### Summary of Updates

Additional experiment testing glycogen synthase inhibition with MZ-101 which successfully normalized cardiac glycogen levels and improved the CS-Mpc2-/- hearts. Additional change was corrections to glucose uptake calculations.

