## Supplementary Fig 1 for "Loss of mitochondrial pyruvate transport initiates cardiac glycogen accumulation and heart failure"

### Supplementary Fig. 1 – Glycolytic pool sizes in failing MPC-/- hearts

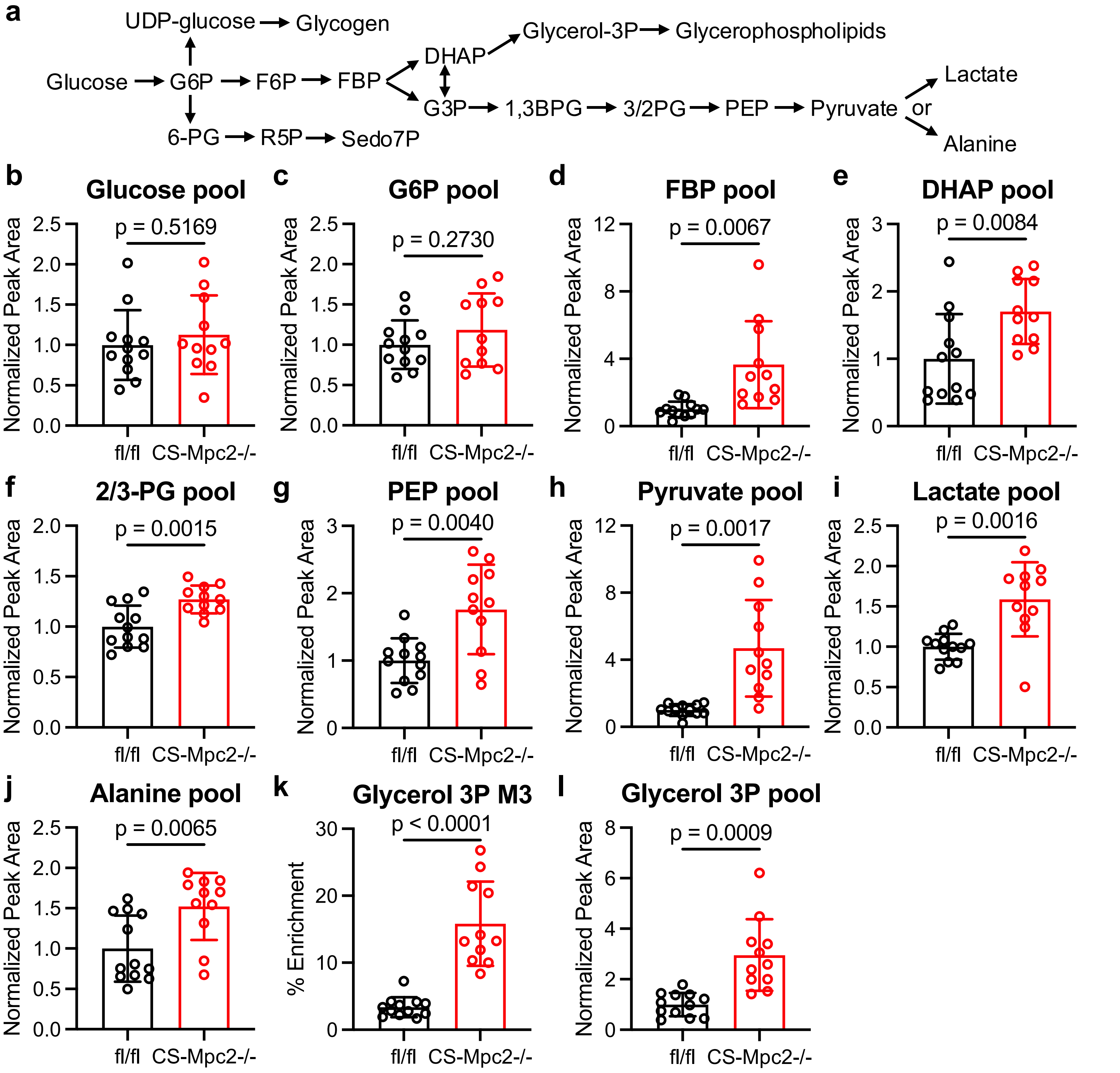


**Supplementary Fig. 1: Glycolytic pool sizes in failing MPC-/- hearts.** Related to Fig. 2. **a** Schematic pathway of glycolysis and accessory glucose pathways. **b-l** Normalized pool size of glucose **(b)**, glucose-6-phosphate (G6P) **(c)**, fructose bisphosphate (FBP) **(d)**, dihydroxyacetone phosphate (DHAP) **(e)**, 2/3-phosphoglycerate (PG) **(f)**, phosphoenolpyruvate (PEP) **(g)**, pyruvate **(h)**, lactate **(i)**, alanine **(j)**, and the $^{13}$C % enrichment of glycerol 3-phosphate (Glycerol 3P) (**k**) and glycerol 3P pool **(l)** of failing CS-Mpc2-/- hearts and fl/fl littermates injected with U-$^{13}$C-glucose, n=11-12. Data are presented as mean±SD. Data were evaluated by unpaired, two-tailed Student's t-test with Welch correction.
