## Supplementary Fig 2 for "Loss of mitochondrial pyruvate transport initiates cardiac glycogen accumulation and heart failure"

### Supplementary Fig. 2 – TCA cycle enrichment and pool sizes in failing hearts

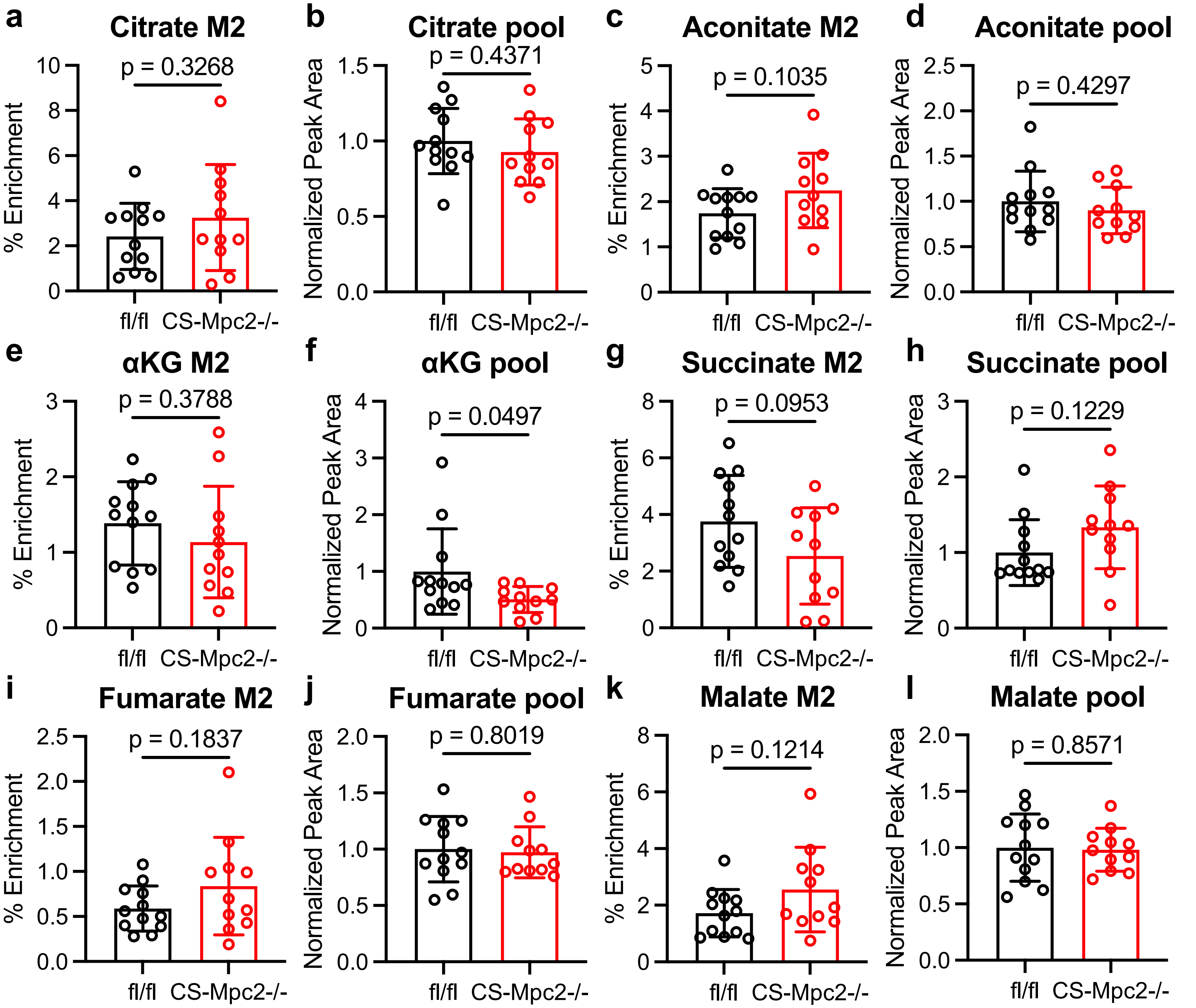


**Supplementary Fig. 2: TCA cycle enrichment and pool sizes in failing CS-Mpc2-/- hearts.** Related to Fig. 2. **a-l** the $^{13}C$ % enrichment and pool sizes of citrate **(a-b)**, aconitate **(c-d)**, $\alpha$-ketoglutarate (KG) **(e-f)**, succinate **(g-h)**, fumarate **(i-j)**, and malate **(k-l)** in fl/fl and failing CS-Mpc2-/- hearts of mice injected i.p. with U-$^{13}C$-glucose, n=11-12. Data are presented as mean±SD. Data were evaluated by unpaired, two-tailed Student's t-test with Welch correction.
