## Supplementary Fig 3 for "Loss of mitochondrial pyruvate transport initiates cardiac glycogen accumulation and heart failure"

### Supplementary Fig. 3 – MPC deletion reduces pyruvate oxidation

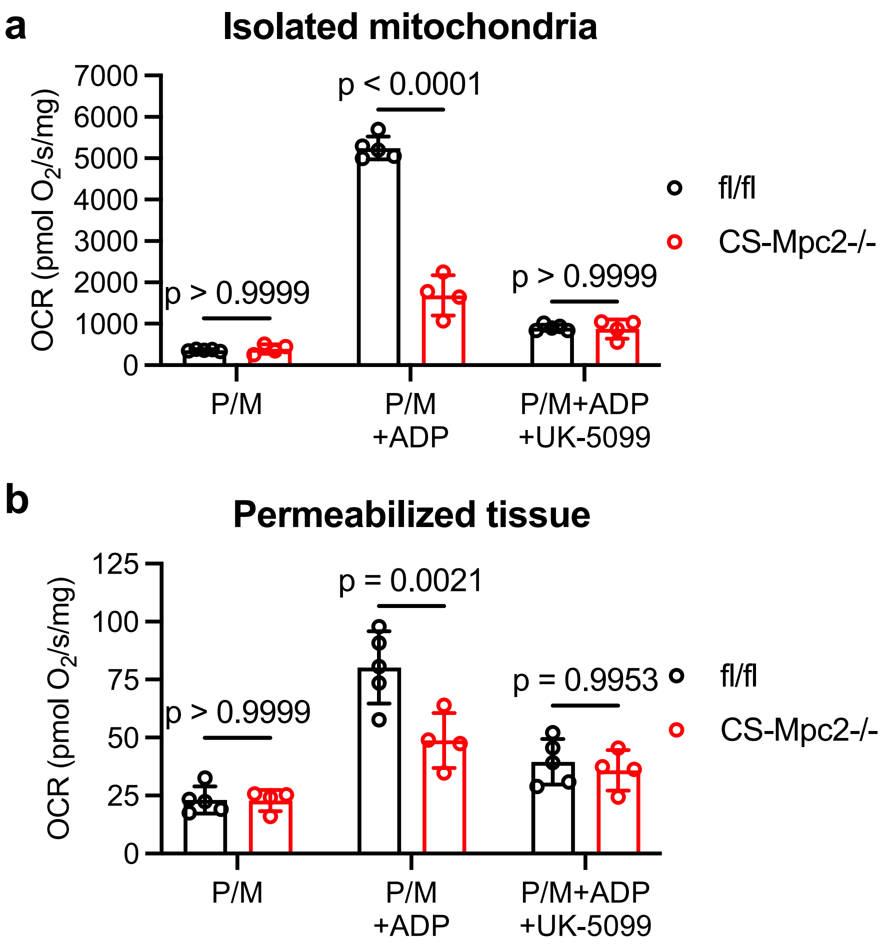


**Supplementary Fig. 3: MPC deletion reduces pyruvate oxidation.** Related to Fig. 2. **a-b** Oxygen consumption rates (OCR) measured from isolated cardiac mitochondria (**a**) and permeabilized cardiac muscle fibers (**b**) from hearts of fl/fl and CS-Mpc2-/- littermates stimulated with pyruvate/malate (P/M), P/M plus adenosine diphosphate (ADP), and P/M, ADP, and the MPC inhibitor UK-5099 (5 μM), n=4-5. Data are presented as mean±SD. Data were evaluated by unpaired, two-tailed Student's t-test with Welch correction.
