## Supplementary Fig 4 for "Loss of mitochondrial pyruvate transport initiates cardiac glycogen accumulation and heart failure"

Supplementary Fig. 4 – No major changes in the pentose phosphate pathway in failing MPC-/- hearts

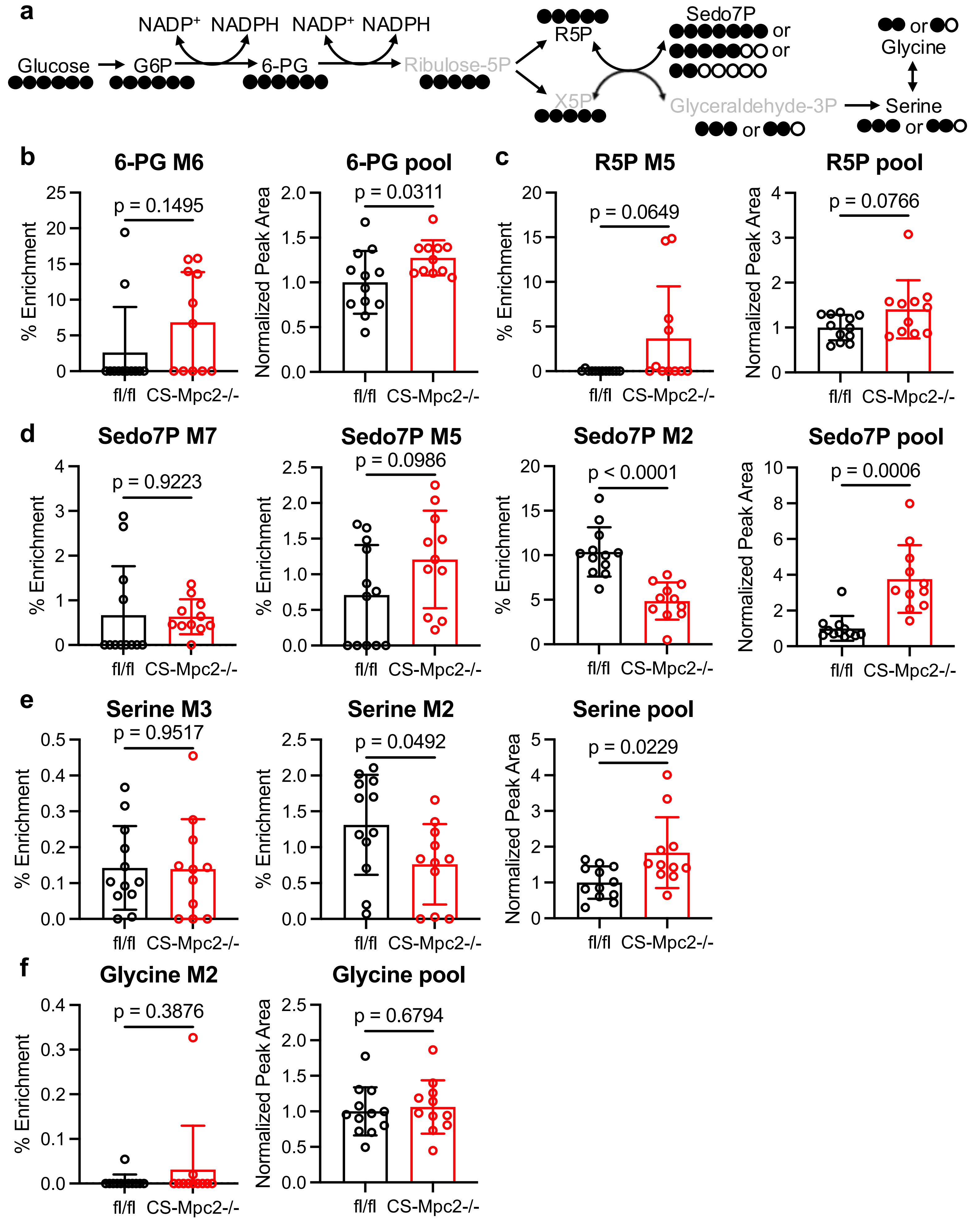

**Supplementary Fig. 4: No major changes in the pentose phosphate pathway in failing MPC-/- hearts.** Related to Fig. 2. **a** Schematic of $^{13}C$ enrichment of the pentose phosphate pathway from U-$^{13}C$-glucose. Metabolites in grey were not measured. **b-f** $^{13}C$ % enrichment and normalized pool size of 6-phosphogluconate (PG) **(b)**, ribose-5-phosphate (R5P) **(c)**, sedoheptulose 7-phosphate (Sedo7P) **(d)**, serine **(e)**, and glycine **(f)**, in hearts from fl/fl and failing CS-Mpc2-/- mice injected with U-$^{13}C$-glucose, n=11-12. Data are presented as mean±SD. Data were evaluated by unpaired, two-tailed Student's t-test with Welch correction.
