## Supplementary Fig 5 for "Loss of mitochondrial pyruvate transport initiates cardiac glycogen accumulation and heart failure"

Supplementary Fig. 5 – Glycolytic pool sizes in young, non-failing MPC-/- hearts

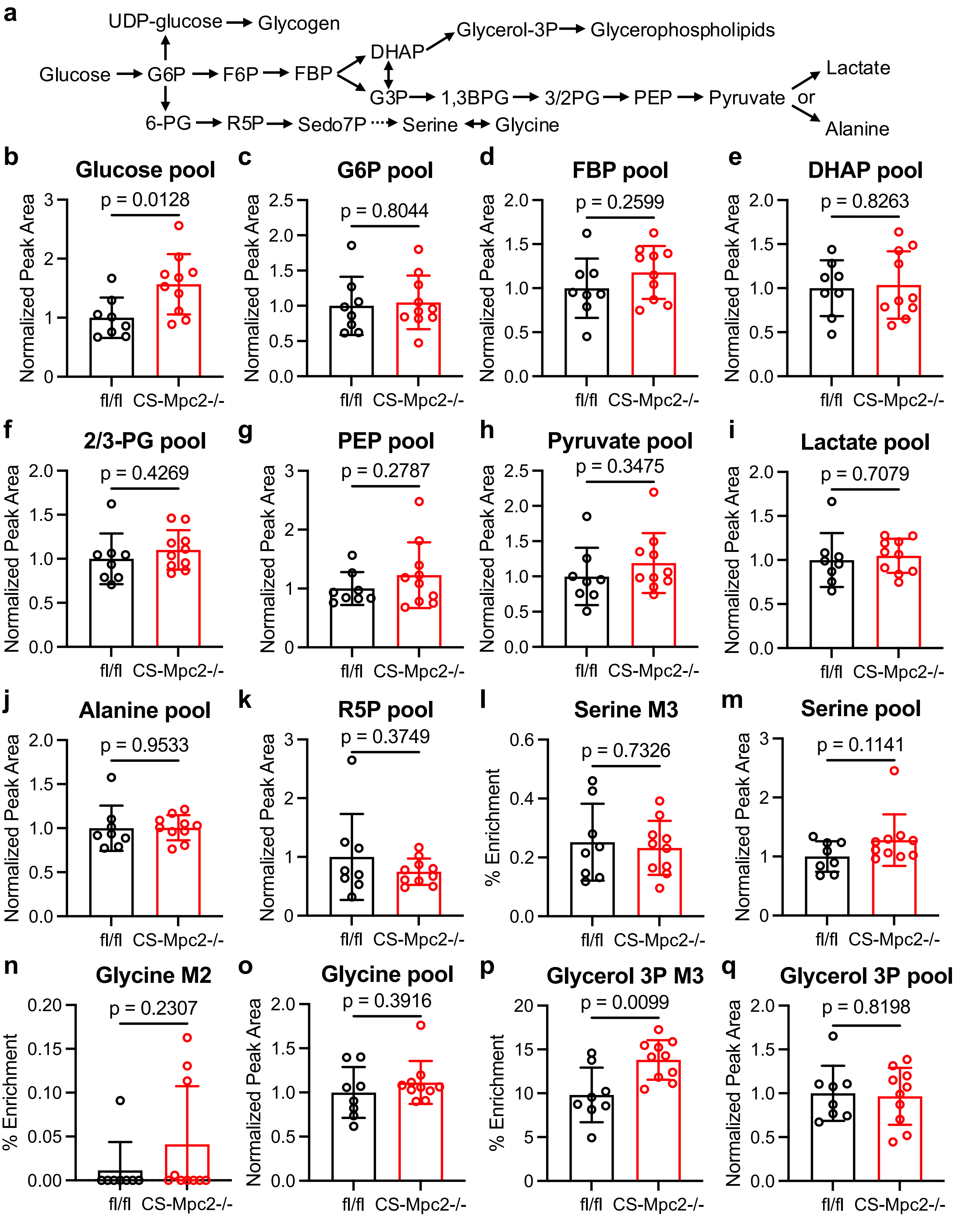

**Supplementary Fig. 5: Glycolytic pool sizes in young, non-failing MPC-/- hearts.** Related to Fig. 4. **a** Schematic of glycolysis and accessory glucose pathways. **b-q** Normalized pool sizes of glucose **(b)**, glucose-6-phosphate (G6P) (**c**), fructose bisphosphate (FBP) (**d**), dihydroxyacetone phosphate (DHAP) **(e)**, 2/3 phosphoglycerate (PG) **(f)**, phosphoenolpyruvate (PEP) **(g)**, pyruvate **(h)**, lactate **(i)**, alanine (j), ribose-5-phosphate (R5P) **(k)**, and the $^{13}C$ enrichment and pool size of serine **(l-m)**, glycine **(n-o)**, and glycerol 3-phosphate (glycerol 3P) **(p-q)** in young non-failing CS-Mpc2-/- and fl/fl littermates injected with U-$^{13}C$-glucose, n=8-10. Data are presented as mean±SD. Data were evaluated by unpaired, two-tailed Student's t-test with Welch correction.
