## Supplementary Fig 6 for "Loss of mitochondrial pyruvate transport initiates cardiac glycogen accumulation and heart failure"

Supplementary Fig. 6 – TCA cycle enrichment and pool sizes in young, non-failing CS-Mpc2-/- hearts

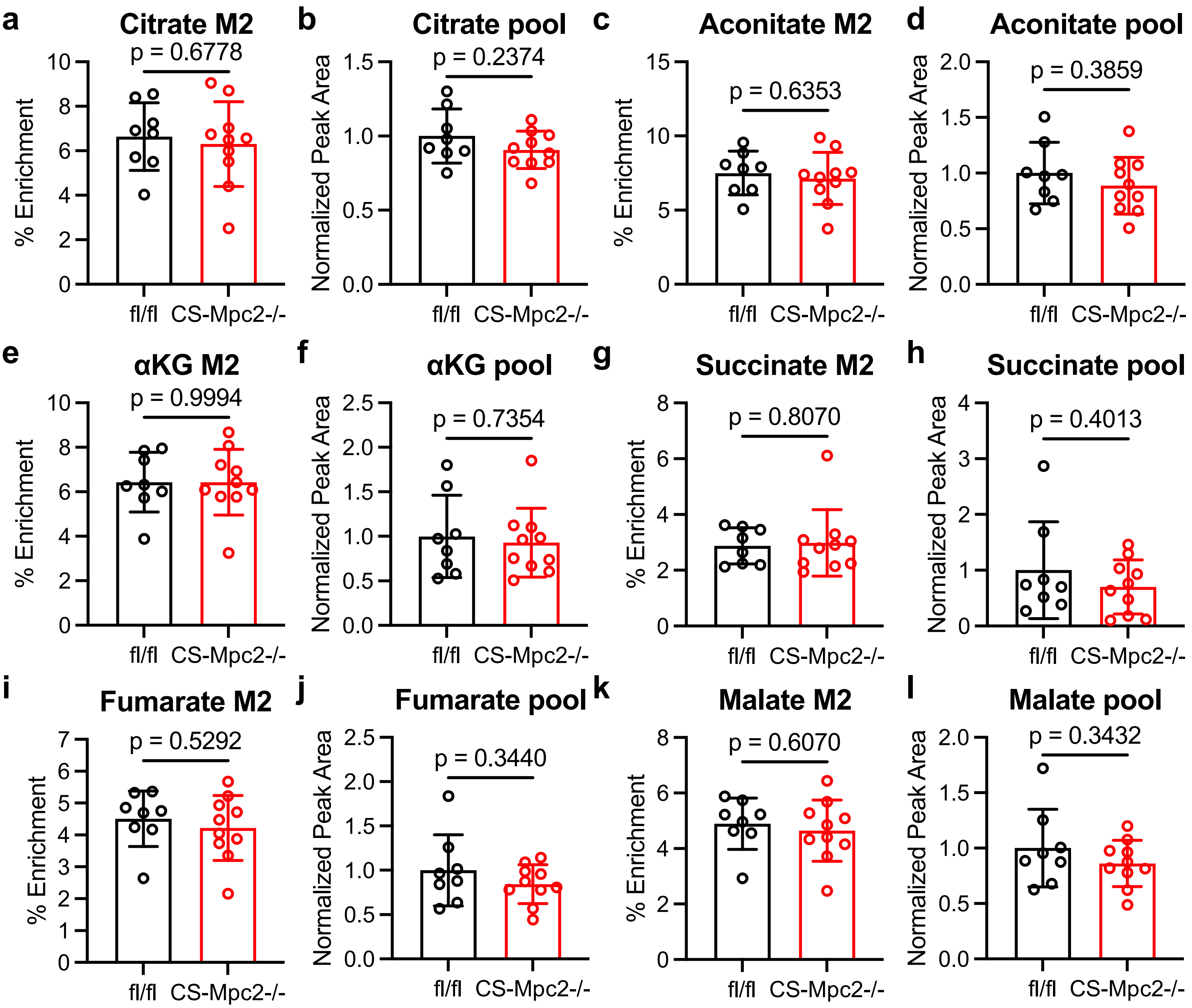


**Supplementary Fig. 6: TCA cycle enrichment and pool sizes in young, non-failing CS-Mpc2-/- hearts.** Related to Fig. 4. **a-l** $^{13}$C % enrichment and normalized pool sizes of citrate (**a-b**), aconitate (**c-d**), α-ketoglutarate (αKG) (**e-f**), succinate (**g-h**), fumarate (**i-j**), and malate (**k-l**), in hearts of young, nonfailing CS-Mpc2-/- and fl/fl littermates injected with U-$^{13}$C-glucose, n=8-10. Data are presented as mean±SD. Data were evaluated by unpaired, two-tailed Student's t-test with Welch correction.
