## Supplementary Fig 7 for "Loss of mitochondrial pyruvate transport initiates cardiac glycogen accumulation and heart failure"

### Supplementary Fig. 7 – Ketogenic diet decreases TCA cycle enrichment from glucose

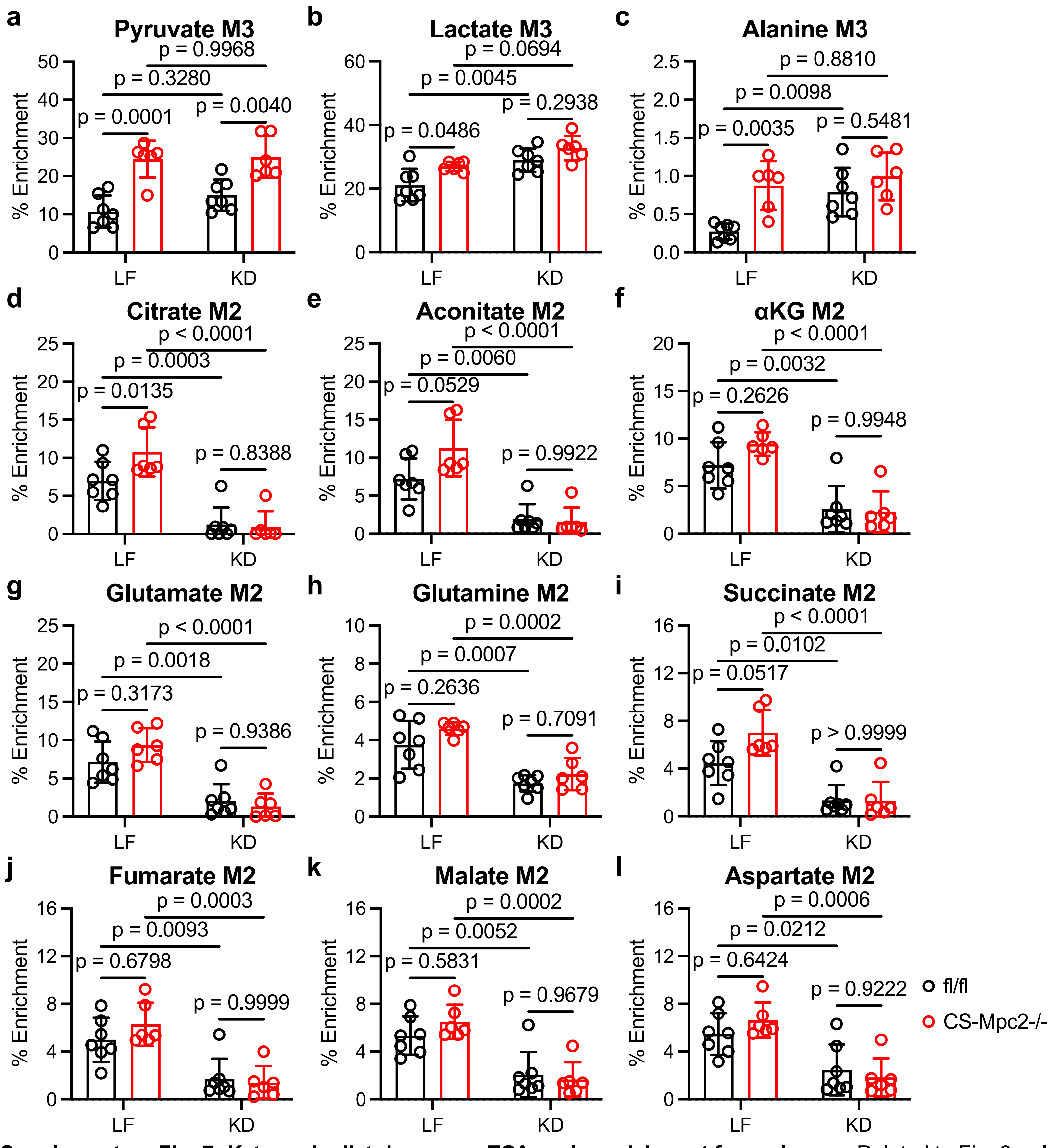


**Supplementary Fig. 7: Ketogenic diet decreases TCA cycle enrichment from glucose.** Related to Fig. 6. **a-l** $^{13}$C % enrichment of pyruvate (**a**), lactate (**b**), alanine (**c**), citrate (**d**), aconitate (**e**), α-ketoglutarate (αKG) (**f**), glutamate (**g**), glutamine (**h**), succinate (**i**), fumarate (**j**), malate (**k**), and aspartate (**l**) in hearts from fl/fl and CS-Mpc2-/- fed either low-fat (LF) or ketogenic diet (KD) and injected with U-$^{13}$C-glucose, n=6-7. Data are presented as mean±SD. Data were evaluated by two-way analysis of variance (ANOVA) with Tukey post-hoc multiple-comparisons test.
